# Re-evaluating functional landscape of the cardiovascular system during development

**DOI:** 10.1101/102723

**Authors:** Norio Takada, Madoka Omae, Fumihiko Sagawa, Neil C. Chi, Satsuki Endo, Satoshi Kozawa, Thomas N. Sato

**Author notes:** Corresponding Author: Thomas N. Sato The Thomas N. Sato BioMEC-X Laboratories Advanced Telecommunications Research Institute International (ATR) 2-2-2 Hikaridai, Seika-cho,Soraku-gun, Kyoto, Japan 619-0288. These two authors contributed equally to this work (Norio Takada, Madoka Omae).

## Abstract

The cardiovascular system facilitates body-wide distribution of oxygen, a vital process for development and survival of virtually all vertebrates. However, zebrafish, a vertebrate model organism, appears to form organs and survive mid-larval periods without the functional cardiovascular system. Despite such dispensability, it is the first organ to develop. Such enigma prompted us to hypothesize yet other cardiovascular functions that are important for developmental and/or physiological processes. Hence, systematic cellular ablations and functional perturbations are performed on zebrafish cardiovascular system to gain comprehensive and body-wide understanding of such functions and to elucidate underlying mechanisms. This approach identifies a set of organ-specific genes, each implicated for important functions. The study also unveils distinct cardiovascular mechanisms, each differentially regulating their expressions in organ-specific and oxygen-independent manners. Such mechanisms are mediated by organ-vessel interactions, circulation-dependent signals, and circulation-independent beating-heart-derived signals. Hence, a comprehensive and body-wide functional landscape of the cardiovascular system reported herein may provide a clue as to why it is the first organ to develop. Furthermore, the dataset herein could serve as a resource for the study of organ development and function.

**SUMMARY STATEMENT:** The body-wide landscape of the cardiovascular functions during development is reported. Such landscape may provide a clue as to why the cardiovascular system is the first organ to develop.

## INTRODUCTION

The cardiovascular system has evolved to facilitate oxygen transport throughout the body (Aaronson et al., 2014; Gabella, 1995). Availability of oxygen is required for development and function of virtually all organs. Oxygen deficiency, referred to as hypoxia, results in developmental and functional failure and/or damages of organs (Semenza, 2011, 2014; Simon and Keith, 2008). Hence, the cardiovascular system is the first functional organ to develop.

The cardiovascular system is also essential for the delivery of humoral factors and immune cells to various parts of the body (Aaronson et al., 2014; Gabella, 1995). Hormones synthesized and secreted by endocrine organs enter into the circulation and reach to other distant organs, where they control organ growth and homeostasis. Immune cells exploit the vascular system to reach to distant tissues where timely immune responses are required. Such responses could facilitate the repair of tissue damages and/or the elimination of intruders and/or unwanted cells such as dead cells and cancer cells. On the other hand, persistence of such responses could result in chronic diseases.

In addition to these canonical functions, the cardiovascular system also locally regulates organ development and function. Hemodynamic force plays important roles in arteriovenous-specification (Sato, 2013), vascular endothelial cell (vEC) behavior (Gimbrone, 1999), vascular extracellular matrix (ECM) remodeling (Califano and Reinhart-King, 2010; Kohn et al., 2015; Reinhart-King et al., 2008), vascular mural cell (vMC) recruitment (Sato, 2013), tissue/organ regeneration (Rabbany et al., 2013) and organ morphogenesis (Serluca et al., 2002). While these locally-acting non-canonical functions are also important, it is generally perceived that the canonical function (i.e., the distantly-acting function) is more predominantly critical for organismal development, function and survival.

Contrary to this perceived predominant importance of the canonical function, zebrafish forms various organs without the functional cardiovascular system (Field et al., 2003a; Field et al., 2003b). Zebrafish is a vertebrate animal, thus with closed circulatory system composed of the heart and the extensive body-wide vascular network. Virtually all vertebrates fail to develop organs without the preceding formation of the cardiovascular system. This is presumably due to the fact that oxygen homeostasis established and maintained by the cardiovascular system is essential for the formation and growth of all functional organs. However, zebrafish appears to form organs even in the absence of the functional cardiovascular system (Field et al., 2003a; Field et al., 2003b). It is assumed that oxygen diffusion through the body wall is sufficient for the initial organogenesis (Field et al., 2003a; Field et al., 2003b). Despite such dispensability for oxygen homeostasis, the cardiovascular system develops first. This observation raises a possibility that the cardiovascular system plays other critical roles in regulating developmental and/or physiological processes. Such cardiovascular function may not be apparent by anatomical observations or the measurements of conventional physiological parameters.

Hence, we re-evaluated the functions of zebrafish cardiovascular system during organogenesis. Specifically, we addressed two questions: 1) What else other than oxygen homeostasis is regulated by the cardiovascular system?; 2) How is it regulated? Experiments were designed to gain comprehensive and body-wide insights into these questions. This was achieved by selectively eliminating each functional aspect of the cardiovascular system in zebrafish larvae. Genetic and pharmacological manipulations were applied to zebrafish larvae to induce ablation(s) and/or functional perturbations of cardiomyocytes, cardiac contraction, the circulation (hemodynamic force), oxygen supply, and/or vECs (vessels)/hematopoietic cells. Molecular signatures induced by each of such manipulations were characterized by body-wide gene expression patterns.

We find the expression of many organ-specific genes are differentially influenced by each manipulation. Such genes include those which are implicated for their functional importance in metabolism, sterol homeostasis, sensory system development and function, and neural functions. Surprisingly, body-wide hypoxia has very small, if any, effects on their expressions. Instead, other distinct cardiovascular mechanisms appear to regulate them. Such mechanisms are mediated by 1) local organ-vessel interactions, 2) circulation-dependent signals, and 3) circulation-independent but distantly-acting beating-heart-derived signals. Hence, these functions are acting more dominantly than the regulation of oxygen-homeostasis during organogenesis. Therefore, such functions could explain why the cardiovascular system develops first. These results also suggest such less-appreciated functions of the cardiovascular system are more important than previously perceived. The findings also provide a body-wide landscape depicting organ-specific gene expression patterns that are differentially regulated by the distinct cardiovascular functions and mechanisms. Such landscape may serve as a resource and a platform for the future in-depth analyses of organ development and function.

## RESULTS

### Characterization of “heartless”

To investigate the role of the heart during development, the zebrafish larvae lacking the heart (referred to as “heartless” here) was characterized. The “heartless” was generated by treating *Tg*(*cmlc2:mcherry-NTR*) zebrafish larvae with MTZ (Curado et al., 2007; Curado et al., 2008; Dickover et al., 2013) (Fig. 1A, see METHODS). The specific ablation of cardiac muscle, but not endocardium/endothelial cells, was confirmed by the lack of cardiac muscle fluorescent reporter expression (*cmlc2:mcherry*) (Fig. 1B), and the presence of endothelial reporter gene expression (*fli1a:egfp*) (Isogai et al., 2003; Lawson and Weinstein, 2002) (Fig. 1B). The lack of circulation was also confirmed (Movies 1, 2). Blood cells were often stuck and accumulated within the heart and between the heart and liver, instead of circulating (Movies 1, 2).

**Figure 1.**
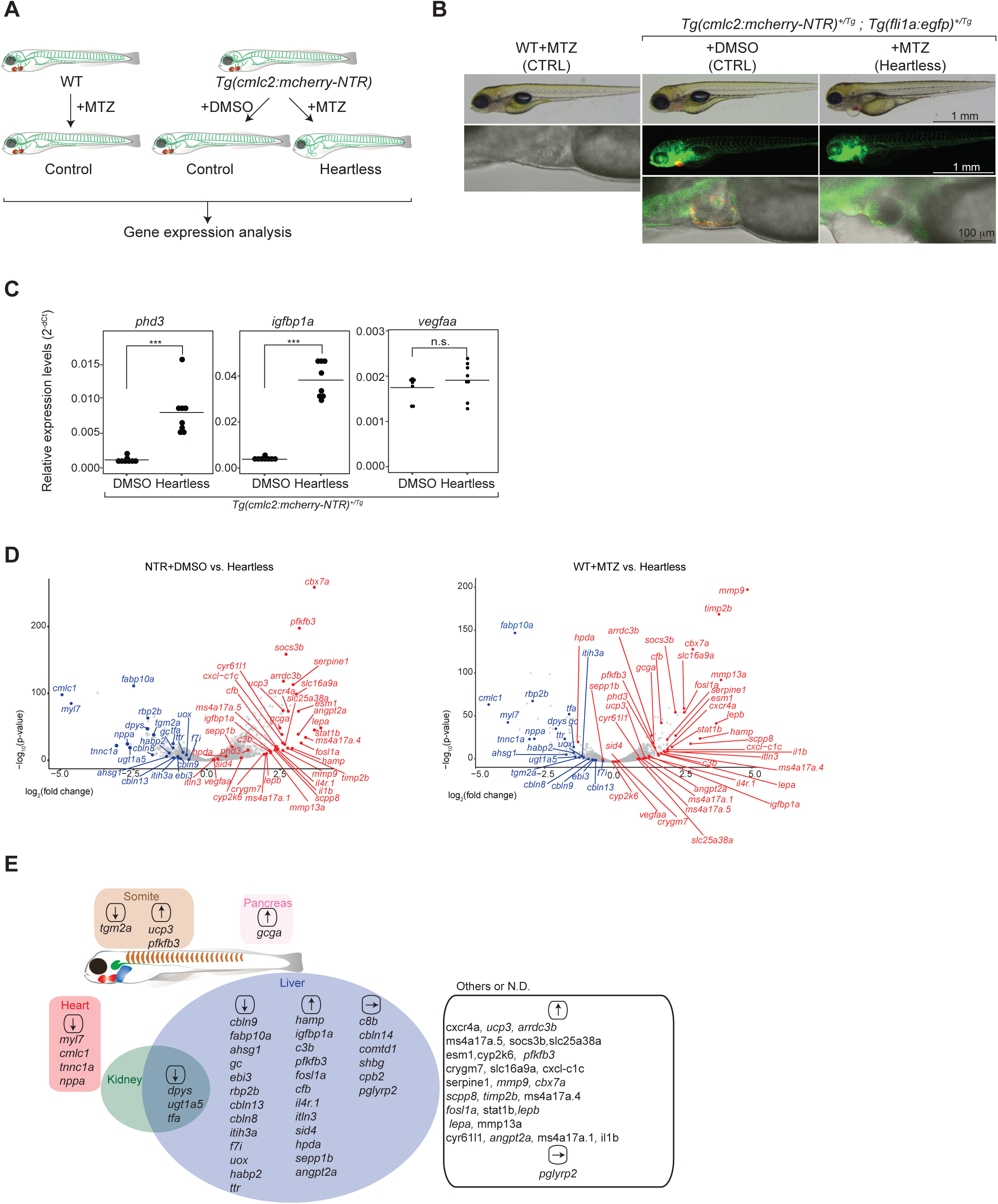
Characterization of “heartless”. **A.** Schematic description of the “heartless” model and the experimental strategy. **B.** Specific absence of cardiomyocytes in “heartless”. Scale bars: 1 mm (top panel), 1 mm (middle panels), 100 μm (bottom panels). Red: cardiomyocyte. Green: vascular endothelial-cells (ECs) and endocardium. Bright-field/DIC images are shown for WT-MTZ as they do not carry a fluorescent reporter. **C.** qRT-PCR analysis of pan-hypoxia indicator genes in “heartless”. ***p<0.001, n.s.: not significant. n=8 for both *Tg*(*cmlc2:mcherry-NTR*)+DMSO and “heartless”. **D.** Volcano plot representing the differentially expressed genes in “heartless”, as compared to *Tg*(*cmlc2:mcherry-NTR*)+DMSO (NTR+DMSO) (left panel) and WT+MTZ (right panel). n=2. Up- and down-regulated genes (as also confirmed by qRT-PCR which is shown in Table S1) are highlighted by red and blue, respectively. **E.** Organ-specific expression patterns of differentially expressed genes in “heartless”. They are placed in each organ. Up- and down-regulated genes are indicated by ↑ and ↓, respectively. Some of the genes unaffected in “heartless” are also included and indicated by →. Many of the liver-specific genes were also confirmed by whole-body transcriptome analysis of the liver-ablated larvae (“liverless”) (Fig. S3).

The state of hypoxia in “heartless” larva was characterized by the expression levels of pan-hypoxia indicators, phd3 (Manchenkov et al., 2015; Santhakumar et al., 2012), igfbp1a(Kajimura et al., 2005) and vegfaa (Wu et al., 2015) that were previously used with zebrafish (Fig. 1C). The expressions of phd3 and igfbp1a were significantly upregulated (Fig. 1C). No significant changes in the vegfaa expression was detected (Fig. 1C), which may reflect a fact that vegfaa is expressed in cardiomyocytes (Zhu et al., 2017). The “heartless” larvae died at around 8 dpf (days post fertilization).

The body-wide gene expression pattern of 4.5 dpf “heartless” was characterized by genome-wide transcriptome analysis. It was compared to two controls, *Tg*(*cmlc2:mcherry-NTR*) treated with the vehicle (DMSO) (NTR+DMSO) and wild type treated with MTZ (WT+MTZ). Differentially expressed genes are shown as Volcano plots (Fig. 1D). The sites of their differential expression were determined by whole-mount in situ hybridization (WISH) analyses (Figs. S1, S2). The results are summarized in Fig. 1E, depicting a body-wide landscape of differentially expressed genes in “heartless”. Many of such genes are primarily expressed in the liver. Furthermore, their functions are implicated for a broad range of biological processes. For example, fabp10a, rbp2b and gc are involved in metabolism of lipids and vitamins. Several genes, such as c3b, cfb, hamp, angpt2a, il4r.1, are implicated for their roles in tissue remodeling and immune responses. This may suggest that liver development and/or function is more dominantly influenced by the cardiovascular system, as compared to those of other organs, at least during this early-to mid-organogenesis periods. The dominant effect on the liver may also suggest the earlier functional maturation of this organ.

### Characterization of cardiomyocyte-specific gene mutations

The MTZ/NTR-mediated cell-ablation system induces cell death of the targeted cells (Curado et al., 2007; Curado et al., 2008; Dickover et al., 2013), thus inflammation could locally occur at the cell-ablated tissues. In fact, some of the genes influenced in “heartless” are related to tissue inflammation (e.g., hamp, il4r.1, il1b, cxcr4a, etc.). Hence, it is important to examine a possibility of an effect of tissue inflammation. Furthermore, the differential gene expression pattern could be due to the lack of cardiomyocytes or cardiac contraction, as both are absent in “heartless”. Thus, these points were addressed by characterizing mutant zebrafish larvae for three cardiomyocyte-specific genes, myl7, cmlc1, tnnc1a, each encoding protein critical for cardiac contraction (Figs. 2A, S4A, S4B, see METHODS). Time-lapse observations of each mutant larva show differential effects on the contraction properties (Movies S3 – S5). The myl7 mutant shows complete lack of contraction (Movie S3). The cmlc1 mutant heart exhibits shuddering movement (Movie S4). The tnnc1a mutant shows significantly reduced ventricular contraction, but relatively normal atrial contraction (Movie S5). These aberrant cardiac contractions also resulted in perturbed circulation (Movies S3 – S5).

The expression analyses of pan-hypoxia indicators show significant upregulation of both phd3 and igfbp1a (Fig. 2B). The expression of vegfaa is significantly upregulated in both cmlc1 and tnnc1a mutants (Fig. 2B), but not in myl7 mutant (Fig. 2B). These results may reflect differences in the degree and/or quality of hypoxic states resulting from the differential cardiac dysfunctions among the mutants.

**Figure 2.**
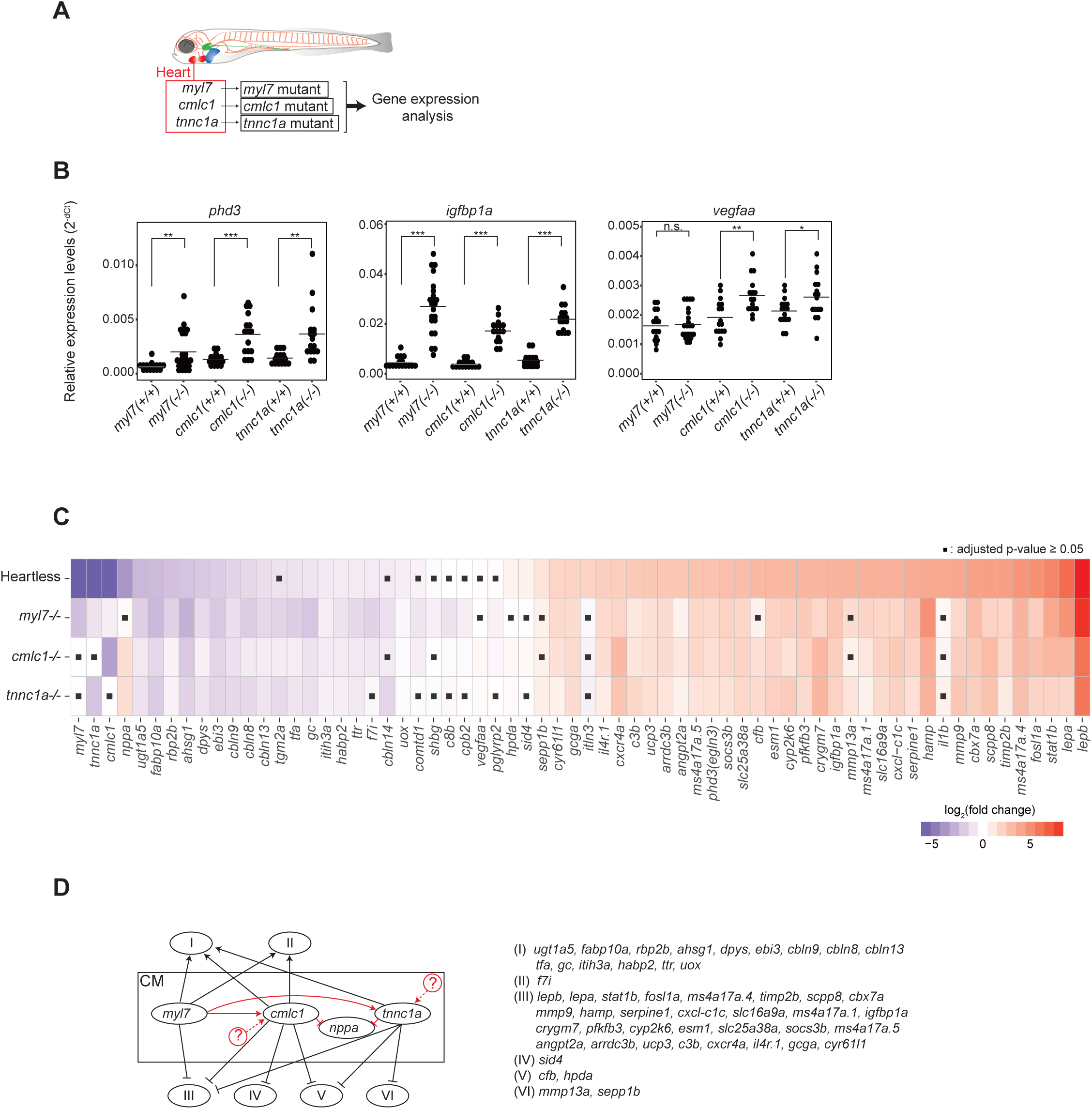
Characterization of cardiomyocyte-specific gene mutants. **A.** Schematic description of the cardiomyocyte-specific gene mutant models and the experimental strategy. **B.** qRT-PCR analysis of pan-hypoxia indicator genes in each mutant. *p<0.05, **p<0.01, ***p<0.001, n.s.: not significant, student-t test. n=16 (WT siblings), 22 (myl7^-/-^), 16 (cmlc1^-/-^), 16 (tnnc1a^-/-^). **C.** Heatmap representing the expression patterns of each gene in each mutant. ▪p≧0.05, student t-test followed by Benjamini-Hochberg procedure to correct errors for the multiple tests. n=16 - 22. **D.** Genetic interactions among mutants. See the text for the detailed description of each category. Black arrows and lines indicate interactions between cardiomyocytes and non-cardiac organs. Red arrows and lines indicate intra-cardiac interactions. A possibility of yet unidentified upstream regulators of cmlc1 and/or tnnc1a are indicated by 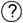

The gene expression patterns of these mutant larvae and “heartless” were compared and summarized (Fig. 2C). The heatmap shows the similar gene expression patterns among “heartless” and the three mutants, despite the lack of apparent cellular ablations in the mutant hearts and variable hypoxic states in the mutants. These results indicate that defective cardiac contraction and/or circulation, but not the local tissue inflammation or the cardiomyocyte-absence itself or hypoxia, is a primary cause of the differential gene expression patterns in “heartless”.

The impact of the myl7 mutation on the gene expression pattern was completely offset (R^2^=0.95501) by the re-expression of myl7 using its own cardiomyocyte-specific promoter (Fig. S4C, D, Table S1). Such gene expression rescue was accompanied by the complete rescue of the normal contractility and circulation (Movie S6). This result further indicates the importance of cardiac contraction to establish and/or maintain the normal body-wide gene expression landscape. The myl7 promoter-driven cardiomyocyte-specific re-expression of cmlc1 or tnnc1a was not sufficient to rescue the defective cardiac contractions or circulation in each corresponding mutant (Fig. S4C, Movies S7, S8, Table S1). Such incomplete functional rescues resulted in only partial or the lack of gene expression rescues (Fig. S4D). These results further support the indication that the normal cardiac contraction and circulation determines the body-wide gene expression landscape.

A few differences in the gene expression patterns were found among the three mutant larvae (Fig. 2C). Such differences could be explained by the presence of genetic interactions between cardiac and non-cardiac genes (black arrows and lines in Fig. 2D). According to the interactions, each non-cardiac gene is categorized into six groups (I – VI) (Fig. 2D): (I) Positive regulation by all three cardiac-genes; (II) Positive regulation by myl7 and cmlc1, but not by tnnc1a; (III) Negative regulation by all three cardiac genes; (IV) Negative regulation by cmlc1, but not by myl7 or tnnc1a; (IV) (V) Negative regulation by cmlc1 and tnnc1a, but not by myl7; (VI) Negative regulation by tnnc1a, but not by myl7 or cmlc1.

In addition to such cardiac and non-cardiac gene interactions (black arrows and lines in Fig. 2D), there also appears an intra-cardiac interaction (red lines in Fig. 2D). The expression of cardiac nppa is suppressed by cmlc1 and tnnc1a, as indicated by its upregulation by the mutation of either gene (Fig. 2C, D). The expressions of cmlc1 and tnnc1a require myl7, as they are both downregulated by the myl7 mutation (Fig. 2C, D).

These results further indicate a differential sensitivity of each gene expression to the cardiac contraction and/or circulation properties. It is possible that subtle changes in the local mechanical signals may also influence gene expressions in non-cardiac organs and also within the heart.

### Possible cross-regulations among liver genes?

Many genes influenced in “heartless” are liver-genes (Fig. 1E). Hence, a possibility of their cross-regulations was examined by analyzing the mutants for 9 liver genes (Fig. S5A, B). Nonsense-mediated-decay (NMD) was confirmed, except for rbp2b (Fig. S5C). The expression of none of the 54 genes regulated in “heartless” was affected by any of these liver gene mutations (Fig. S5D, Table S1), indicating the existence of very little, if any, one-on-one cross-regulations among these liver-genes.

### Characterization of “vesselless”

Oxygen and humoral factors are delivered to tissues and organs through the circulation. Immune cells use the vessels to reach to their target tissues. The circulation requires cardiac contraction and vessels. Therefore, the absence of either one results in the lack of circulation. Hence, any effects caused by the lack of circulation are likely to be shared between “heartless” and larvae without vessels. In contrast, the gene expression regulated by the local vEC-derived signal(s) could be influenced only in the larvae without vessels, but not in “heartless”. To distinguish between these two types of regulations, we generated the larvae without the vessels but with intact contracting cardiac-muscle. This was made possible by using *cloche/npas4l* mutant (Liao et al., 1997; Reischauer et al., 2016)(referred to as “vesselless” here) (Fig. 3A). The absence of the vessels and endocardium, but the presence of cardiomyocytes (Fig. 3B) and cardiac contraction (Movie S9), was confirmed.

**Figure 3.**
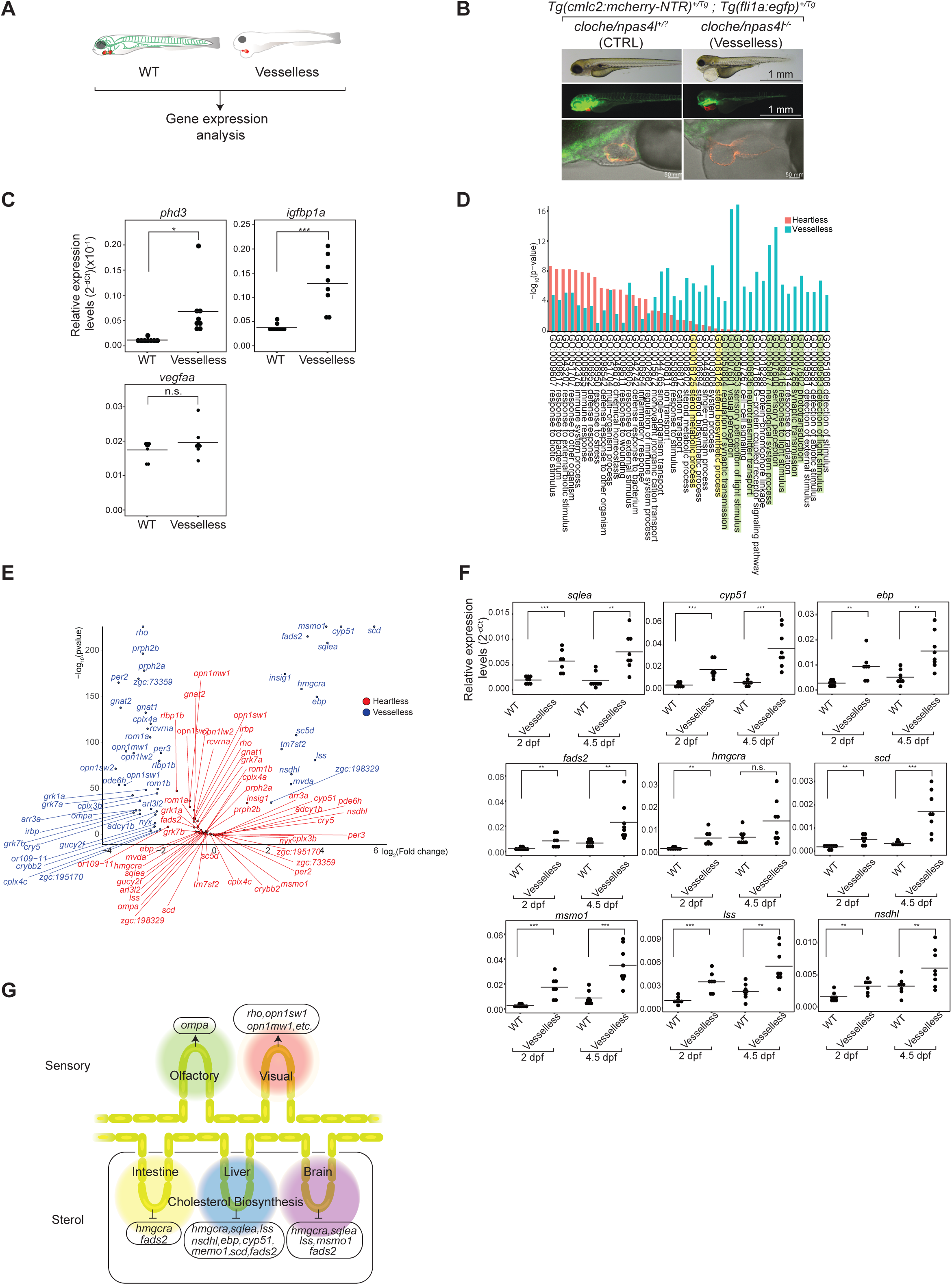
Characterization of “vesselless”. **A.** Schematic description of the “vesselless” model and the experimental strategy. Green: vECs, endocardium and hematopoietic cells. Red: Cardiomyocytes. **B.** Specific absence of hematovascular cells (i.e., vECs, endocardium, hematopoietic cells) in “vesselless”. Green: vECs and endocardium. Red: cardiomyocyte. Scale bars: 1 mm (top panel), 1 mm (middle panels), 50 μm (bottom panels). **C.** qRT-PCR analysis of pan-hypoxia indicator genes in “vesselless”. *p<0.05, ***p<0.001, n.s.: not significant, student-t test. n=8 (WT siblings), 8 (“vesselless”). **D.** GO enrichment analysis of “vesselless” (blue) and “heartless” (red). The top 44 GO terms are shown. Highlighted are neural/sensory (green) and sterol (yellow) related GOs. **E.** Volcano plot representing differentially expressed genes in “vesslless”. Shown are genes that are specifically affected in “heartless” (red) and in “vesselless” (blue). n=2. **F.** qRT-PCR analysis of cholesterol biosynthesis genes. **p<0.01, ***p<0.001, n.s.: not significant, student-t test. n=8 (WT siblings), 7 (“vesselless” at 2dpf), 8 (“vesselless” at 4.5dpf). **G.** Summary of differentially expressed genes in “vesselless”.

The upregulation of two hypoxia indicators, phd3 and igfbp1a, was detected in “vesselless”, as in “heartless” (Fig. 3C). No significant changes of vegfaa expression was detected in “vesselless”, like in “heartless” (Fig. 3C), which may in part reflect a possibility of its expression in hematopoietic cells (Liao et al., 1997; Reischauer et al., 2016). These results indicate that the degree and the quality of body-wide hypoxia in “vesselless” is comparable to that in “heartless”. Hence, it is likely that the difference between “vesselless” and “heartless” is the absence/presence of the vessels, but not the states of the cardiac-contraction or hypoxia.

The comprehensive transcriptome analysis of 5 dpf “vesselless” was conducted to identify genes whose expressions are specifically dependent on the presence of the vessels (Figs. 1B, 3B). The GO enrichment analysis unveiled families of genes whose expressions are specifically affected in “vesselless”, but not in “heartless” (Fig. 3D, E). They include those related to sensory/neural system (i.e., synaptic transmission, response to light stimulus, sensory perception, phototransduction, detection of light stimulus, neurological system process, neurotransmitter transport, sensory perception of light stimulus, visual perception, regulation of synaptic transmission) and sterol homeostasis (i.e., sterol biosynthetic process, sterol metabolic process) (Fig. 3D, E). They are affected only in “vesselless”, but not in “heartless”, despite hypoxia and the lack of circulation in both.

Many sensory/neural system genes are specifically downregulated in “vesselless” (Fig. 3E). One such gene is olfactory-marker-protein-a (ompa) (Fig. 3E). WISH analyses show ompa is expressed in the olfactory bulb (Fig. S6). A comparison to a pan-olfactory-bulb marker, ompb (Celik et al., 2002; Yoshida et al., 2002; Yoshida and Mishina, 2003), shows that both ompa and ompb are expressed in the olfactory bulb, but the ompa expression is restricted to a sub-domain of the neuroepithelium (Fig. S6).

Several sterol homeostasis-related genes are specifically upregulated in “vesselless” (Fig. 3E). They include a family of genes encoding enzymes critically involved in cholesterol biosynthesis (Lu et al., 2015; Mazein et al., 2013; Paton and Ntambi, 2009) (Fig. S7). WISH analyses indicate that all were expressed in the liver, and five (hmgcra, sqlea, lss, msmo1, fads2) were also expressed in the brain and two (hmgcra, fads2) in the intestine (Fig. S8). Cholesterol in the circulation is taken up by vECs via endocytosis (Anderson et al., 2011; Ho et al., 2004). This vEC mechanism is critical to maintain the cholesterol level in circulation (Anderson et al., 2011; Ho et al., 2004). It is possible that this mechanism operates as a negative feedback, suppressing the expression of the genes encoding cholesterol biosynthesis enzymes – hence, the absence of vECs in “vesselless” could induce their upregulation.

This possibility was further supported by an experiment using atorvastatin, a potent inhibitor of HMG-CoA reductase, an enzyme required for cholesterol biosynthesis (D’Amico et al., 2007; Ho et al., 2004). The atorvastatin treatment of wild type or “heartless” zebrafish larvae induced the upregulation of these genes (Fig. S9, Table S1). In contrast, the atorvastatin treatment of “vesselless” induced only a weak or no upregulation (Fig. S9, Table S1).

We also examined their expression in 2 dpf larvae where major organs, such as liver, are yet to grow, but vECs are already present, thus an indirect pleiotropic effect is minimized (Fig. 3F). The result shows that their expressions are also upregulated in 2 dpf “vesselless” larvae (Fig. 3F). These results suggest that their expressions are in a negative feedback loop where vECs function as a suppressive interface.

That such differential expressions of the genes were due to the lack of vessels is further supported by the characterization of etv2/etsrp morphant (Fig. S10). Etv2/etsrp morpholino injection was previously shown to reduce the vascular network in a relatively specific manner (Craig et al., 2015; Sumanas and Lin, 2006; Veldman and Lin, 2012) (Fig. S10A). Gene expression correlation analysis between “vesselless” and etv2/etsrp morphant showed a high correlation coefficient (R^2^=0.9379582) (Fig. S10B, Table S1), supporting the indication that the differential gene expressions in “vesselless” is due to the lack of vessels, rather than a pleiotropic effect of the *cloche/npas4l* gene mutation.

Taken together, the results suggest that vECs themselves function as positive and negative regulators for sensory gene and sterol homeostasis gene expressions, respectively (Fig. 3H). In “vesselless”, the absence of vEC-derived signals, such as cell-cell contacts, ECM, secreted paracrine factors, may induce the downregulation of sensory system genes such as ompa and opsin/rhodopsin genes in olfactory and visual systems, respectively (Fig. 3E, G). The absence of vECs in “vesselless” also induces the lack of cholesterol endocytosis, causing the elimination of the negative feedback loop of cholesterol biosynthesis gene expression and hence the upregulation of their expression (Fig. 3E, F, G).

### Distinguishing differential roles of the cardiovascular system in regulating the body-wide gene expression landscape

The “heartless” lacks cardiomyocytes, resulting in the absence of cardiac contraction and circulation (Fig. 1B, Movie S2). The cardiac gene mutants exhibit no or perturbed cardiac contractions and show the lack of circulation (Movies S3 – S5). In “vesselless” larvae, cardiomyocytes and contraction are present (Fig. 3B, Movie S9), but lacks the vessels (Fig. 3B), thus resulting in the lack of circulation (Movie S9). The tissues in “heartless”, cardiac-gene mutants and “vesselless” are under hypoxia (Figs. 1C, 2B, 3C). Hence, some, if not all, of the differential gene expressions detected among these models could be caused by one or the combinations of differential conditions of each model, but not by the common ones such as hypoxia. To determine which of these conditions are contributing to the differential gene expression patterns, we examined additional models.

One such is a cardiotoxin-treated model (Fig. 4A). Haloperidol is known to disturb the normal cardiac contraction in zebrafish (Milan et al., 2006). While cardiomyocytes and vasculature were present as in normal larvae (Fig. 4B), the zebrafish larvae treated with this drug for 5 hrs exhibited arrested heart-beat and no circulation (Movie S10). Hypoxia of the haloperidol-treated larvae was characterized by the expression of pan-hypoxia indicators (Fig. 4C). The haloperidol-treatment caused significant upregulated expression of all three pan-hypoxia indictors (Fig. 4C).

**Figure 4.**
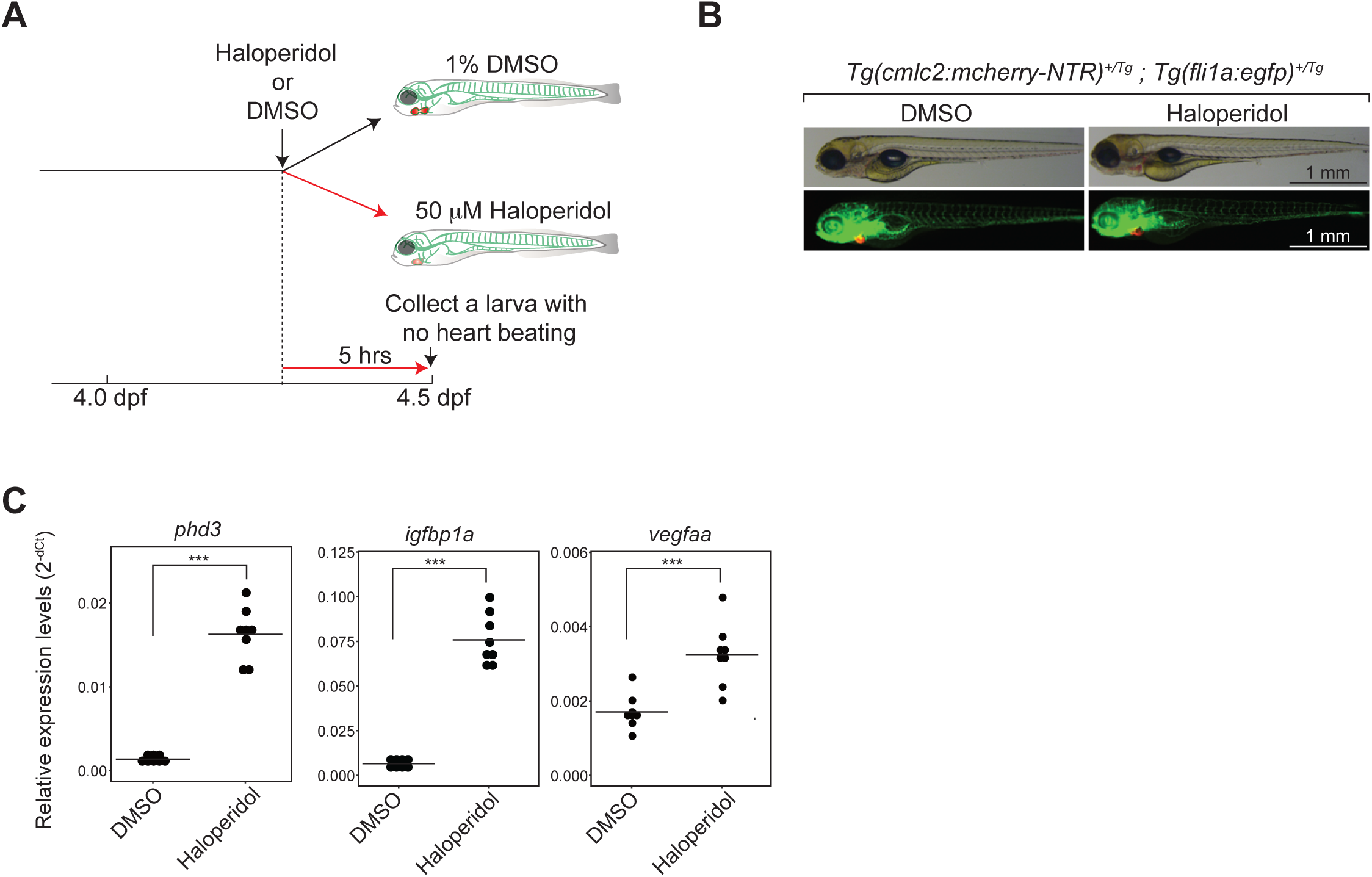
Characterization of a cardiotoxin-treated model. **A.** Schematic description of a cardiotoxin-treated model and the experimental strategy. **B.** Presence of cardiomyocytes and hematovascular cells in the model. Green: hematovascular cells; Red: cardiomyocytes. Scale bar, 1 mm. **C.** qRT-PCR analysis of pan-hypoxia indicator genes. ***p<0.001, student-t test. n=8 (DMSO treated control), 8 (Haloperidol-treated).

Contributions of hypoxia were also examined (Fig. 5). The body-wide hypoxia was induced by two methods (Fig. 5A): DMOG-treatment and hypoxia-chamber (Gerri et al., 2017). The larvae were treated with 100 μM DMOG for 6 hrs, 10 hrs, or with 125 μM for 24 hrs (Fig. 5A, see also METHODS). The heart-beat and the circulation appeared relatively normal in the DMOG-treated larvae (Movie S11). Body-wide hypoxia was also induced by incubating the larvae in hypoxia-chamber for 24 hrs (3.5 dpf – 4.5 dpf) (Fig. 5A, see also METHODS). The 24-hrs treatment/incubation period in the hypoxia models are comparable to the duration of the cardiac-ablation in “heartless” (see METHODS). Hypoxia was evaluated by the expressions of pan-hypoxia indicators (Fig. 5B). In both models, the expression of phd3 was significantly upregulated (Fig. 5B). The expressions of both ifgbp1a and vegfaa expression were upregulated in the larvae treated by DMOG for 10 hrs and 24 hrs, and also in those incubated in hypoxia-chamber for 24 hrs (Fig. 5B).

**Figure 5.**
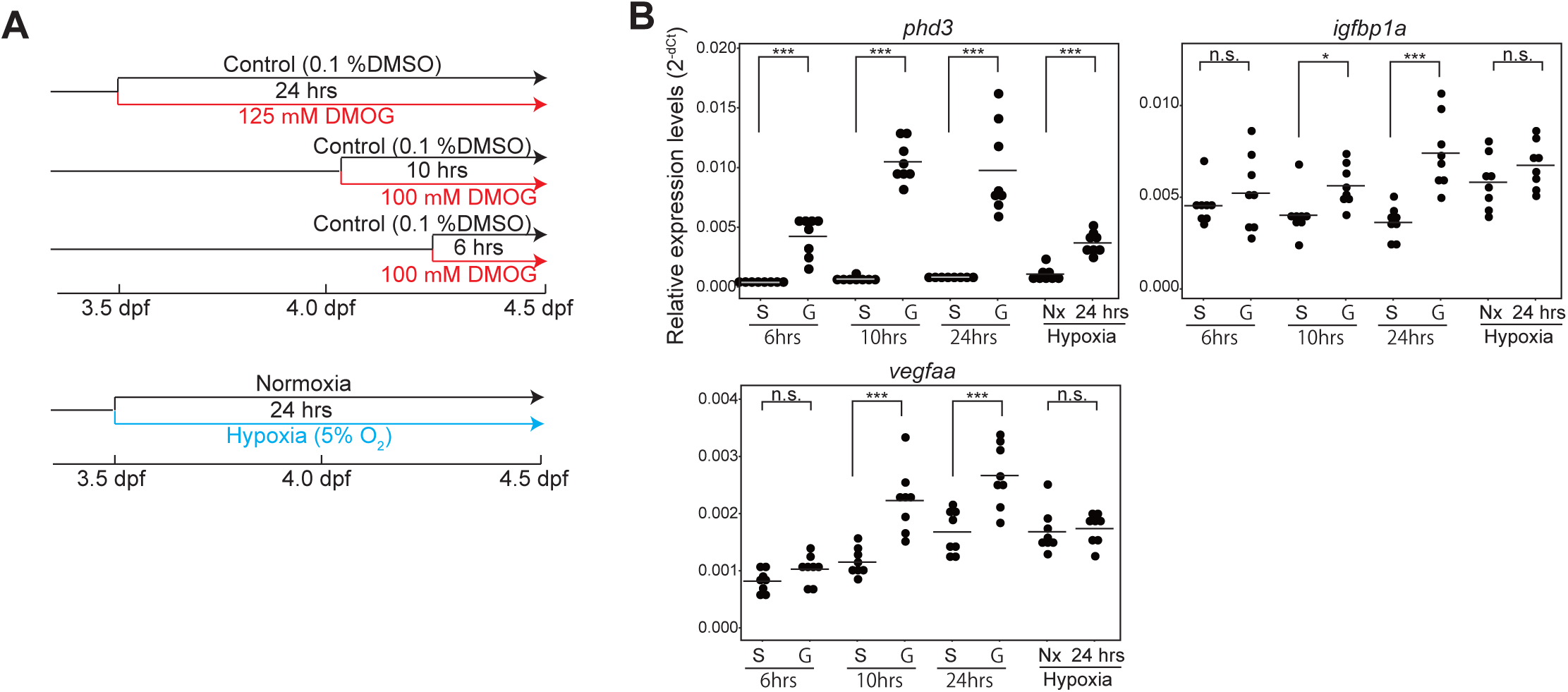
Characterization of hypoxia models. **A.** Schematic description of hypoxia models and the experimental strategy. **B.** qRT-PCR analysis of pan-hypoxia indicator genes. *p<0.05, ***p<0.001, n.s.: not significant, student-t test. n=8 (DMSO treated control), 8 (6 hrs DMOG-treated), 8 (10 hrs DMOG-treated), 8 (24 hrs DMOG-treated), 8 (24 hrs normoxia control), 8 (24 hrs hypoxia chamber).

The differential gene expression patterns were analyzed in these additional models and compared to those of “heartless” and “vesselless” (Fig. 6). We also introduced the vessel-ablation to “heartless” (i.e., “heartless+vesselless”) to determine the dependence of the differential gene expression in “heartless” on the presence of vessels (Fig. 6). The structural and functional characteristics of each model are summarized in Fig. 6A. The differential gene expressions among the models are summarized and presented as heatmap (Fig. 6B) with the original raw data in Table S1.

The heatmap shows that many of the differential gene expressions found in both or either “heartless” and/or “vesselless” are unaffected in the hypoxia models (i.e., DMOG and hypoxia-chamber). This result indicates that the hypoxic condition induced up to 24 hrs of DMOG or hypoxia-chamber incubation is less influential on the expressions of many of these differentially regulated genes in “heartless” and/or “vesselless”. Based on the degree of the upregulation of phd3, a pan-hypoxia marker, a comparable degree of hypoxia appears to be achieved by “heartless”, “vesselless”, “heartless+vesselless”, DMOG and hypoxia-chamber models (Fig. 6B).

**Figure 6.**
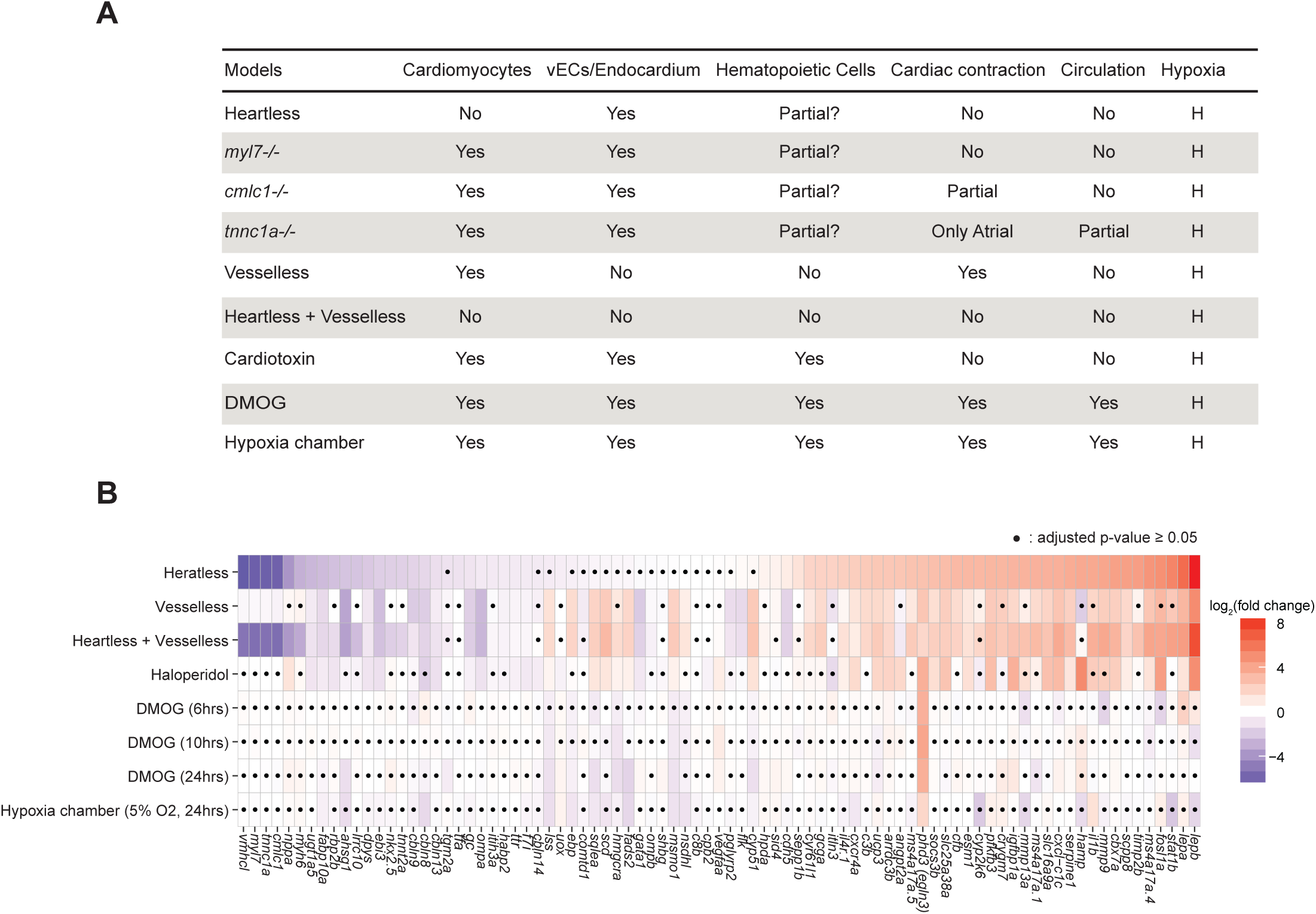
Comparison of the models. **A.** Summary of the characteristics of each model. **B.** Heatmap representation of differential gene expression in each model. ▪p≧0.05, student t-test followed by Benjamini-Hochberg procedure to correct errors for the multiple tests. n=8.

We identified 8 genes that show significantly downregulated expression in “heartless”, “vesselless”, “heartless+vesselless” and haloperidol models, but not in neither of the hypoxia models (i.e., DMOG and hypoxia-chambers) (Fig. 6B). They are ugt1a5, dpys, ebi3, cbln13, gc, ompa, ttr and f7i (Fig. 6B). We also identified 10 genes that show significantly upregulated expression in “heartless”, “vesselless”, “heartless+vesselless” and haloperidol models, but not in neither of the hypoxia models (i.e., DMOG and hypoxia chambers) (Fig. 6B). They are lepb, lepa, ms4a17a.4, scpp8, slc16a9a, pfkfb3, esm1, slc25a38a, ucp3 and il4r.1 (Fig. 6B). The result indicates that the circulation contributes to the regulations of these gene expressions (8 downregulated and 10 upregulated genes). Such circulation-dependent signals could be mechanical, humoral and/or cellular signals. The result also suggests that oxygen homeostasis contributes to much lesser extent, if any, to their expressions.

The expression of ompa is significantly downregulated in “vesselless”, but very little in “heartless” (Figs. 3E, G, 6B). While a small downregulation of ompa is found in “heartless” (log_2_fold=-0.86, p=3.48x10^-6^) and in the haloperidol (log_2_fold=-1.07, p=1.85x10^-5^) models, more robust downregulation (log_2_fold=-2.66, p=4.03x10^-10^) was detected in “vesselless” (Fig. 6B, Table S1). No influences were detected in either of the hypoxia models (Fig. 6B. Table S1). These results suggest that the local-presence of vECs themselves in olfactory bulb is a main contributor to the regulation of the ompa expression. The circulation appears to influence to much lesser extent, and the hypoxia impose little, if any, influence on its expression.

The genes encoding a family of cholesterol biosynthesis enzymes (lss, ebp, sqlea, scd, hmgcra, fads2, msmo1, nsdhl, cyp51) are all significantly upregulated in “vesselless”, but not in “heartless” (Figs. 3F, 6B, Table S1). Much less upregulation was found for some of them in the haloperidol-treated larvae (Fig. 6B, Table S1). Furthermore, in neither of the hypoxia models, their upregulations were found. In fact, only small downregulation was detected for some of them (Fig. 6B, Table S1). Such results suggest that their expressions are mainly regulated by the local presence/absence of vECs, rather than the circulation or hypoxia.

Retinol binding protein 2b (rbp2b) exhibits a distinct expression pattern (Fig. 6B). Rbp2b is implicated for vitamin A and lipid homeostasis (Kanai et al., 1968; Li and Norris, 1996) and primarily expressed in the liver (Figs. S1, S3). The expression of rbp2b is significantly downregulated in “heartless”, but not in “vesselless” (Fig. 6B). Such downregulation in “heartless” is maintained in the “heartless+veseelless” combination model (Fig. 6B, Table S1). Furthermore, the significant downregulation was also detected in the haloperidol model (Fig. 6B). These results suggest that the liver expression of rbp2b requires cardiac contraction, but the vessels are dispensable.

What could be such a mediator relayed from the beating heart to the liver in the absence of the vessels? A possible candidate is a nervous-system-derived mediator. The innervation of the cardiomyocytes is critical for the cardiac function. Dopaminergic (DA) neurons are known to regulate a number of hemostatic processes involving cardiovascular functions (Aaronson et al., 2014; Myers and Olson, 2012; Noble, 2002). Hence, we examined a role of DA neurons in regulating the liver rbp2b expression. A subset of DA neurons was genetically ablated by the double mutations of otpa and otpb genes (Fernandes et al., 2013) (Fig. S11A, B). The ablation of the otpa^+^/otpb^+^ DA neurons was confirmed by the suppression of oxt expression as previously described (Fernandes et al., 2013) (Fig. S11C). However, the rbp2b expression was not affected in the otpa/otpb double mutants (Fig. S11C). Such dispensability of otpa^+^/otpb^+^ DA neurons suggests an existence of other type(s) of beating-heart derived vessel-independent signals in regulating the liver rbp2b expression.

## DISCUSSION

It has been assumed that the cardiovascular system in zebrafish is dispensable for oxygen homeostasis at least during early-to mid-larval periods. This notion suggests an existence of other important cardiovascular function(s) during development. Herein we show several evidences suggesting that vascular-organ interactions, circulation-dependent signals, and circulation-independent but distantly acting beating-heart-derived signals are important mediators of such non-oxygen regulating functions (Fig. 7).

**Figure 7.**
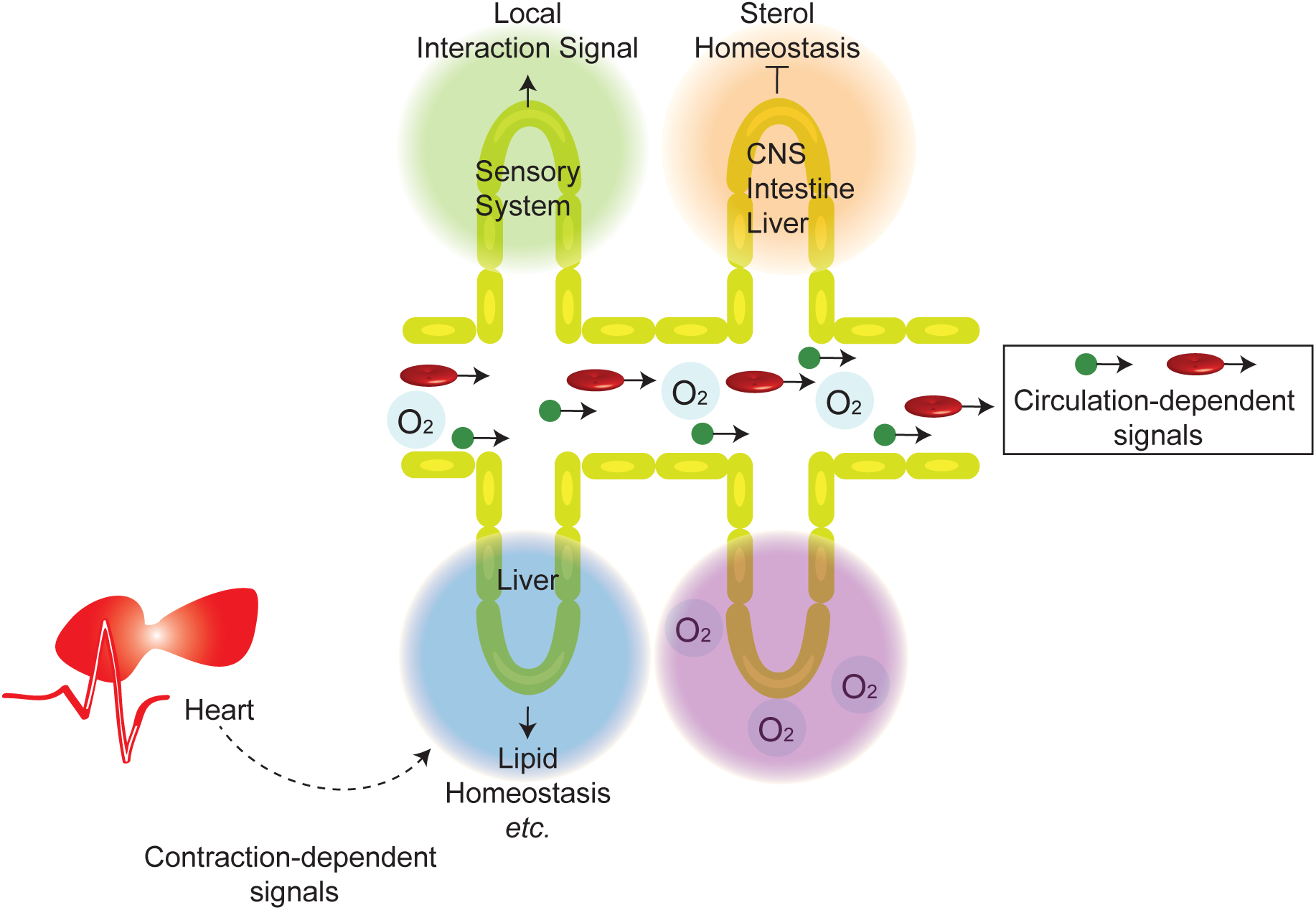
Distinct organ-specific roles of the cardiovascular system during development. See DISCUSSION for the description.

What could be the identities of these signals? For the sensory system (e.g., the ompa expression in the olfactory bulb), vEC-derived signals such as cell-cell contacts, ECM and/or secreted paracrine factors are potential candidates (Crivellato et al., 2007). Although such vEC-derived signals are the main regulators, the circulation also appears to influence the ompa expression (Fig. 6B, Table S1). Therefore, it is possible that the strength/levels of such vEC-derived signals may be partially dependent on the circulation. Alternatively, the mechanical signals mediated by the circulation could function together with the vEC-derived signals to regulate the expression of the sensory system genes. In the case of locally regulating the sterol homeostasis genes by vECs, a negative-feedback loop system, such as cholesterol endocytosis, installed within vECs is a potential candidate (Anderson et al., 2011; Ho et al., 2004).

We also found 18 genes that are regulated by circulation-dependent signals, but not much by hypoxia (Fig. 6B). Mechanical stimuli, such as shear-stress, could be such circulation-dependent signals (Sato, 2013) (Fig. 7). Humoral factors (Sato, 2013) and immune cells and/or factors presented by them are also candidates.

A beating-heart-derived long-distance acting signal is another one. An example regulated by such a signal is rbp2b (Fig. 7). We show that otpa^+^/otpb^+^ DA neurons are dispensable for this regulation (Fig. S11C). What could be such a signal? A possibility of diffusible small chemicals and/or peptides regulated and released by the contracting cardiac-muscle (Fig. 7) would be worth exploring in the future.

We also show that each of such signals and cardiovascular functions are highly selective for specific organs (Fig. 7). In particular, the local vascular-organ interactions appear to be preferentially exploited by sensory organs (such as olfactory and visual systems) (Figs. 3G, 7). Our results also indicate a critical importance of the vascular interface for maintaining cholesterol homeostasis in brain, liver and intestine. In this case, the vasculature appears to function as suppressive interface to prevent hyper-activation of the cholesterol biosynthesis pathways (Figs. 3G, 7). It is possible that the usage of such local cardiovascular functions is more effective for certain organs (e.g., sensory organs) than others. Alternatively, such special organs may lack a system that can utilize circulation-dependent signals or may be less efficient in using them.

Our finding also has an evolutionary implication. We identified a possibility that the heart, via contraction, sends signal(s) to the liver in a circulation-independent manner, as indicated by the differential expression patterns of rbp2b (Figs. 6B, 7). This mechanism could be related to a primordial function of the cardiovascular system. Like in vertebrates, the heart is the first functional organ system to develop also in invertebrates. However, invertebrates lack extensive vascular network. The beating-heart alone is sufficient to facilitate the body-wide transport of oxygen and humoral factors. Hence, it is possible that such signaling system by the beating-heart without the use of the vessel has survived natural selection and has remained in some vertebrates such as zebrafish.

Do our findings apply to other vertebrates such as mammals? In mice and frog, the critical importance of local interactions between vECs and developing organs have been shown for the development of liver and pancreas, respectively (Cleaver and Dor, 2012; Lammert et al., 2001, 2003; Talavera-Adame and Dafoe, 2015). Recently, an importance of vEC-tissue interactions is also implicated in liver organoid formation in vitro (Camp et al., 2017). Neurovascular interactions play critical roles in development and disease in mice and human (Crivellato et al., 2007; Lawson et al., 2002; Mukouyama et al., 2002; Okabe et al., 2014; Visconti et al., 2002; Wang et al., 1998; Weinstein, 2005). Several paracrine factors, collectively referred to as angiocrine factors, have been discovered (Butler et al., 2010a; Ding et al., 2010; Ding et al., 2011). They are secreted from tissue-specific vECs and facilitate organ regeneration (Butler et al., 2010b; Rafii et al., 2016). These findings illustrate an importance of such local-functions of the vasculature in other vertebrates including mammals and human.

Our results also indicate that the zebrafish larvae without the functional cardiovascular system (i.e., “heartless” and “vesselless”) are under hypoxia at least based on the upregulated expression of two pan-hypoxia indicators, phd3 and igfbp1a (Fig. 6B). Previously, it was assumed that zebrafish without the functional cardiovascular system can form organs as oxygen diffusion through body wall is sufficient at least up to mid-larval period. However, our results indicate that the functional cardiovascular system is indeed necessary for oxygen homeostasis and zebrafish organs form under hypoxic microenvironment.

Experimental manipulations of the cardiovascular system induce changes in oxygen-homeostasis, making it challenging to study the roles of the cardiovascular system in regulating oxygen-independent developmental and/or physiological processes. Here, we identified a set of non-cardiovascular genes (e.g., ompa, fabp10a, dpys, ugt1a5, pfkfb3, etc.) that are regulated by the formation and functions of the cardiovascular system, but are only little, if any, influenced by oxygen homeostasis. Hence, these non-cardiovascular genes could be utilized as the targets of manipulations to determine biological processes that are independent of oxygen homeostasis. No direct manipulations of the cardiovascular system are required in this approach, thus it is applicable to mammalian models. For example, the inhibitions of such gene functions could identify biological processes that are independent of oxygen homeostasis, but are mediated by the cardiovascular system and are critical for organismal development and homeostasis, in mammalian models such as mice.

The dataset presented herein also provides a list of marker genes for distinct functional aspects of the cardiovascular system. These markers are useful for evaluating functional manipulations of the cardiovascular system in the future experiments. Furthermore, many diseases are caused by the changes of the functional states of the cardiovascular system (Noble, 2002). Hence, it is also possible that the differential gene expression patterns reported here could be exploited to evaluate the effects of therapeutic treatments on the cardiovascular system in diseases and/or disease models.

In addition, the dataset could also serve as a useful resource to design experiments to gain further in-depth insights into the roles of the cardiovascular system in regulating organ development and function. The olfactory vasculature could function as a guide to induce the differentiation of a ompa-positive subset of neuroepithelial cells. Such vasculature-guided ompa-positive neuroepitheial cells could possess a unique physiological function. In each organ, only subsets of genes are responsive to the manipulations of the cardiovascular system. These cardiovascular-sensitive genes may collectively assume unique developmental and/or physiological functions during organ maturation. Such questions could be systematically addressed by using the annotations of the genes reported in this paper.

## METHODS

### Fish husbandry

Zebrafish were maintained in circulation-type aquarium system (Iwaki) with 14 h/day and 10 h/night cycle at around 27°C. The fertilized eggs were collected and raised at 28.5°C in egg water (0.06% artificial marine salt supplemented with 0.0002% methylene blue) until around epiboly stage and subsequently in 1/3 Ringer’s medium (1.67 mM HEPES, 38.7 mM NaCl, 0.97 mM KCl, 0.6 mM CaCl_2_, pH 7.2) containing 0.001% phenylthiourea (PTU) (Sigma) to prevent pigmentation. Embryos and larvae were staged to days post fertilization (dpf) according to Kimmel *et al* (Kimmel et al., 1995). Zebrafish maintenance and experiments were conducted in accordance with animal protocols approved by the Animal Care and Use Committee of Advanced Telecommunications Research Institute International (A1403, A1503, A1603).

### Transgenic reporter fish

The following zebrafish lines were used: *Tg(cmlc2:mcherry-NTR)*(Dickover et al., 2013), *Tg(fabp10a:CFP-NTR)* (Curado et al., 2007; Curado et al., 2008), *Tg(fli1a:egfp)*^*y1*^(Lawson and Weinstein, 2002) and *Tg(gata1:DsRed)* (Traver et al., 2003).

### Cardiomyocyte / liver ablation

The cardiomyocyte-specific ablation was performed by treating *Tg(cmlc2:mcherry-NTR)* by Metronidazole (MTZ). *Tg(cmlc2:mcherry-NTR)* heterozygous fish were crossed with wild-type to obtain *Tg(cmlc2:mcherry-NTR)*^*+/Tg*^ eggs. Eggs were raised at 28.5°C in egg water to around epiboly stage and subsequently in 1/3 Ringer’s medium containing 0.001% PTU to prevent pigmentation. At 2 - 3 dpf, the embryos expressing the cardiac mcherry fluorescence reporter were selected under Leica M165 FC microscope (Leica). At 3 dpf, the embryos were treated either with 0.2% DMSO alone and 10 mM MTZ with 0.2% DMSO in 1/3 Ringer’s media containing 0.001% PTU for 6 hrs, followed by washing with 1/3 Ringer’s medium 3 times for 5 min each. After the washing, embryos were again raised in 1/3 Ringer’s medium containing 0.001% PTU for 20 hrs at 28.5°C.

For the liver-specific ablation, *Tg(fabp10a:CFP-NTR)* homozygous fish were used. *Tg (fabp10a:CFP-NTR)* embryos/larvae were treated with 7 mM MTZ prepared as above from 2.5 dpf to 5.5 dpf MTZ-containing media was changed daily during the treatment.

### Cloche/npas4l mutant

*Cloche*^*la1164*^*(clo*^*la1164*^) (Liao et al., 1997; Reischauer et al., 2016) (provided by Dr. Kawakami) was maintained by mating with wild-type fish. Genotyping of *clo*^*la1164/la1164*^ was conducted by observing the heart morphology and the absence of the circulation of embryo. C*lo*^*+/la1164*^ was crossed with *Tg(cmlc2:mcherry-NTR)*^*+/Tg*^ and *Tg(fli1a:egfp)*^*+/y*^ to generate double heterozygous fish of *clo*^*la1164*^ *allele* and a reporter gene allele. To genetically ablate the cardiomyocyte in *clo*^*la1164/la1164*^ embryo, *clo*^*+/la1164*^ was mated with *clo*^*+/la1164*^*;Tg(cmlc2:mcherry-NTR)*^*+/Tg*^. At 3 dpf, *clo*^*+/?*^;*Tg(cmlc2:mcherry-NTR)*^*+/Tg*^ and *clo* ^*la1164/la1164*^;*Tg(cmlc2:mcherry-NTR)*^*+/Tg*^ were treated with 0.2% DMSO and 10 mM MTZ in 1/3 Ringer’s medium with 0.001% PTU/0.2% DMSO for 11 hrs. After washing out the solution, the embryos were raised as described in the cardiomyocyte-ablation section.

### RNA extraction

To obtain total RNA, embryos and larvae were harvested in a 1.5 - 2 ml tube at appropriate stage and frozen in liquid N_2_ to be stored in -80°C. To prepare total RNA for RNAseq analysis of embryonic and larval stages, 10 - 20 embryos and larvae were pooled in a 1.5 ml tube, and total RNA was isolated using RNeasy Mini Kit (QIAGEN). The pooled embryos/larvae were homogenized in Buffer RLT included in the kit using 5 ml syringe and 24G needle by passing through the needle for 20 times. Alternatively, they were crashed in 700 μl Buffer RLT using approximately 50 zirconia balls with 1.5 mm diameter (YTZ balls) (NIKKATO) by centrifuging at 4260 rpm for 60 sec in Cell Destroyer PS1000 (BMS). After homogenization, the manufacture instruction was followed. To prepare RNA for real-time PCR analysis, embryos/larvae were individually harvested in a 1.5 ml or 2 ml tube and total RNA from each individual embryo/larva was isolated by AllPrep DNA/RNA Mini Kit (QIAGEN). Individual Embryo/larva was either homogenized using a syringe as described above or crashed in 700 μl Buffer RLT using approximately 50 zirconia balls with 1.5 mm diameter (YTZ balls) (NIKKATO) by centrifuging at 4260 rpm for 60 sec in Cell Destroyer PS1000 (BMS). After the homogenization or crushing, the manufacture instructions were followed. Subsequently, the isolated genomic DNA and total RNA were subjected to genotyping and reverse transcription reaction.

### RNA sequencing

Total RNA was prepared from two biological replicate pools of 4.5 dpf wild type and cardiomyocyte-ablated larvae and 5.5 dpf wild type, cloche mutant and the liver-ablated larvae. Each pool had about 15 larvae. The RNAseq analyses (read length:100 bp, total reads number per sample: about 100 million, single-end read) for Figs. 1 and S3 were performed using Illumina HiSeq 2500. RNAseq analysis (read length: 50 bp, total reads number per sample: about 30 million, single-end read) for WT+MTZ data of Fig. 1 was performed with Illumina HiSeq 1500. The data were mapped to the zebrafish genome (Zv9) with Bowtie2 (Langmead and Salzberg, 2012; Nellore et al., 2016) running on Galaxy (https://usegalaxy.org/) (Afgan et al., 2016). The obtained bam file was used to calculate fragments-per-kilobase-of-exon-per-million (FPKM) of transcripts and the differential gene expression data using Cuffdiff (Trapnell et al., 2010). To perform Gene ontology enrichment analysis, the enriched genes were defined as those with log_2_fold ≥1 and with p <0.05. A gene ontology enrichment analysis is performed by R package “topGO” using a root category “BP” and a reference database “org.Dr.eg.db”. To prepare volcano plot graph from RNAseq data, p-value and fold-change was calculated using DESeq2 (Love et al., 2014) with default settings.

### Quantitative RT-PCR (qRT-PCR) analysis

Total RNA (30 - 150 ng) was used to perform reverse transcription using SuperscriptIII reverse transcriptase primed with oligo(dT) (Invitrogen). After the reaction, the mixture was diluted to 1:3 - 1:10 to prepare a working solution. Real-time qPCR was performed using LightCycler 480 SYBR Green Master (Roche) in combination with LightCycler 480 machine (Roche). The final reaction mixture (10 μl volume) was prepared as followings: 5 μl of LightCycler 480 SYBR Green I Master, 2 μl of RNase-free water, 0.5 μl of 10 μM forward primer, 0.5 μl of 10 μM reverse primer, 2 μl of cDNA template. The mixture was dispensed using epMotion P5073 automated pipetting system (Eppendorf). All qPCR was performed using 384 white-plate PCR platform. PCR cycle was as followings: 10 minutes at 95 °C, 45 cycles of 10 sec. 95°C, 10 sec. at 63°C, 10 sec at 72°C. Amplification and dissociation curves were generated by the LightCycler 480 Software (release 1.5.1.62 SP2). Primers used for qPCR were designed using the Roche Universal ProbeLibrary Assay Design Center (https://qpcr.probefinder.com/organism.jsp). The primer sequences were listed in supplementary table 2. The transcript levels of measured genes were normalized with *rpl13a* level for all experiments.

### Whole-mount in situ hybridization (WISH)

To synthesize an antisense RNA probe, the template DNA was amplified by PCR using KOD-Plus-Neo (TOYOBO) from cDNA synthesized from zebrafish total RNA of appropriate stage of WT or cloche mutant (for hmgcra, sqlea, nsdhl, cbp, fads2). For lss, cyp51, msmo1, scd, ompa and ompb, the cDNA sequences were chemically synthesized for use as templates of PCR. The primers used for PCR are listed in Supplementary Table 1. Primer sequence was designed using Blast primer (NCBI/ Primer-BLAST: http://www.ncbi.nlm.nih.gov/tools/primer-blast/) and, T3 and T7 sequence were added at 5’ end of forward and reverse primer, respectively. PCR product was purified by QIAquick PCR Purification Kit (Qiagen). The sequence was confirmed using T3 and T7 primers. For synthesizing antisense RNA probe, the following mixture was used: DIG RNA labeling mix (Roche Diagnostics), Transcription buffer (Roche Diagnostics), RNase inhibitor (Roche Diagnostics), T7 RNA polymerase (Roche Diagnostics), and 200 ng of template DNA. Then it was incubated for 1.5 - 3 hour at 37°C, followed by precipitation with lithium-chloride precipitation solution. Precipitated DIG-labeled RNA was re-suspended in nuclease free water and mixed with equal volume of formamide to be stored at -80°C.

To prepare embryos for whole mount in situ hybridization, anesthetized zebrafish embryos/larvae were fixed in 4% Paraformaldehyde Phosphate Buffer Solution (PFA) (Nacalai Tesque) overnight at 4°C. The fixed embryos were dehydrated three times for 5 minutes in 100% methanol at room temperature and were stored at -30°C for at least 2 days. Before hybridization, the embryos in methanol were dehydrated five times for 5 minutes in PBST (phosphate buffered saline containing 0.1% Tween-20), and then permeabilized in 10 μg/ml proteinase K in PBST for 30 minutes at room temperature. After quick wash with PBST, the embryos were post-fixed for 20 minutes in 4% PFA at room temperature, and then washed 5 x 5 minutes in PBST. Then, the embryos were incubated in hybridization solution (50% formamide, 5×SSC, 50 μg/ml Heparin, 500 μg/ml tRNA, 0.1% Tween-20, 9.2 mM Citric acid, pH 6.0) without DIG-RNA probe at 68°C for at least one hour. Hybridization was performed in hybridization solution containing DIG-labeled probe (1:200 dilution) at 68°C for 16 hours. Following to hybridization, the embryos were washed with 50% formamide /50% 2×SSC once for 5 minutes and then for 15 min at 68 °C, and then 2 x SSC once for 15 minutes at 68°C, finally followed by the wash with 0.2xSSC twice for 30 minutes at 68°C. After wash with PBST 5 x 5 min, the embryos were blocked with blocking solution of 2% normal sheep serum (NIPPON BIO-TEST LABORATORIES INC.), 2 mg/ml BSA at room temperature for one hour, and then incubated with anti-Digoxigenin-AP Fab fragments (Roche) (1:5000 dilution) in blocking solution for overnight at 4°C. After the incubation, the embryos were washed with PBST five times for 15 minutes at room temperature, and then washed in coloration buffer (50 mM MgCl_2_, 100 mM NaCl, 0.1% Tween20, 100 mM Tris-HCl, pH 9.5) at room temperature. DIG was detected with BM purple AP solution (Roche) at 4°C

When the desired staining intensity was reached, the embryos were washed 3 x 5 minutes with PBST, then fixed in 4% PFA with 0.1% glutaraldehyde (Wako). After fixation, embryos were placed in 75-80% glycerol in PBS.

### Two-color fluorescence in situ hybridization

DNP-labeled RNA probe was synthesized using T7 RNA polymerase (Roche) by incubating template DNA with 0.35 mM DNP-11-UTP (Perkin Elmer), 1 mM of ATP, GTP and CTP, 0.65 mM UTP (Invitrogen) for 3 hrs at 37°C. The synthesized DNP-probe was purified by LiCl precipitation.

Embryos were prepared and hybridized with DIG-labeled and DNP-labeled riboprobe as described in WISH using AP-system, except for addition of 5% dextran sulfate (Sigma) to the hybridization buffer. The hybridized embryos were washed and blocked as WISH method of AP-system. After blocking, to detect DIG-labeled probe, the embryos were incubated with anti-digoxigenin-POD, Fab fragments (1:1000; Roche) in PBST containing 2% sheep serum and 2 mg/ml BSA for overnight at 4°C. After the incubation, the embryos were washed 6 x 15 min in PBST and then 2 x 5 min in 1x amplification diluent, followed by the incubation with TSA Plus Cyanine 5 solution (1:50 dilution in amplification diluent buffer) (Perkin Elmer) for 1 hr at R.T. After the incubation, embryos were washed 2 x 5 min with PBST and then the first TSA reaction was quenched in 2% H_2_O_2_ in PBST for 60 min at R.T. The embryos were then washed 4 x 5 min in PBST, followed by the incubation with anti-DNP-HRP (1:500; Perkin Elmer) in PBST containing 2% sheep serum and 2 mg/ml BSA for overnight at 4°C. The antibody was washed out in PBST, and then the embryos were incubated in TSA Plus Cyanine 3 (1:50 dilution in amplification diluent buffer) (Perkin Elmer) for 1 hr at R.T. After the incubation, the embryos were washed 6 x 5 min in PBST and mounted in Prolong Diamond (Molecular probes) for imaging under confocal microscope.

### Microscopy and image process

To observe and record the heart beating and the circulation, embryos/larvae were anesthetized using 0.012% MS-222 and were mounted either laterally or ventrally in 1.0% NuSive GTG Agarose (Lonza) on glass-bottomed 35 mm dish. Imaging was performed using a 10x dry objective lens (Plan Apo, NA0.45) and 20x dry objective lens (Plan Apo, NA 0.75) mounted on Nikon A1R confocal microscope. Time-lapse image was recorded with a resonant scanner for 15-30 f/s imaging, and converted to Quick time movie using IMARIS software (BITPLANE) or to AVI movie using Figi software. These movies were converted into mp4 movies using iMovie software.

To take images of whole mount in situ hybridization, specimens were mounted in 75-80% glycerol and imaged using 4 x (Plan Apo/NA0.20) or 10 x (Plan Apo/NA 0.45) (Nikon) objective lens mounted on Nikon eclipse inverted microscope and 1 x objective lens (Plan Apo) mounted on Leica M165 FC microscope. Images of two-color fluorescence WISH were taken using a 20x dry objective lens (Plan Apo, NA 0.75) and 40x water immersion objective lens (Apo LWD, NA 1.15) mounted on Nikon A1R confocal microscope.

### CRISPR/CAS9 mutagenesis

For CRISPR/Cas9, sgRNAs were designed using the online tool CHOPCHOP (http://chopchop.cbu.uib.no/#), CRISPR DESIGN (http://crispr.mit.edu/) and CRISPRdirect (https://crispr.dbcls.jp/). Target sequences and guide RNA (gRNA) sequences were listed in Table S2. For preparing gRNA, we followed either plasmid-based method, where the template sequence for gRNA was cloned in plasmid, or cloning-free method. For the plasmid based method(Jao et al., 2013), two complementary 20 μl base oligonucleotides corresponding to the target sequence were annealed in 20 μl solution with 1x NEBuffer3 by the following procedure: denaturation for 5 min at 95°C, cooling to 50°C at -0.1°C /sec, pausing at 50°C for 10 min and cooling to 4°C at -0.1°C /sec. One μl of annealed oligonucleotides was mixed with 400 ng of pT7-gRNA (Jao et al., 2013) (Addgene), which is a gRNA cloning vector, 0.5 μl of three restriction enzymes of BsmBI, BglII and SalI (NEB) each, 1 μl of 10x NEBuffer3 and 1 μl of T4 DNA ligase (NEB) in a volume of 20 μl to perform digestion and ligation in a single step. The oligonucleotides/enzymes mixture was incubated in three cycles of 20 min at 37°C, 15 min at 16°C, followed by 10 min at 37°C and 15 min at 55°C. Two μl of the reaction mixture was used to transform DH5α. After the preparation of plasmid using QIAprep Spin Miniprep Kit (QIAGEN) from several colonies, the successful cloning was confirmed by sequencing with M13Forward primer. The plasmid with gRNA target sequence was linearized by BamHI and used as a template of *in vitro* transcription reaction. gRNA was transcribed using MEGAshortscript kit (Ambion). The cloning-free method was also used to generate templates for gRNA synthesis (Gagnon et al., 2014). The 1 μl of 100 μM gene-specific oligonucleotides containing T7 or SP6 sequence, 20 base target sequence without PAM, and a complementary region to constant oligonucleotide were mixed with 1 μl of 100 μM oligonucleotide encoding the reverse-complement of the tracrRNA tail with 1x NEBuffer2 in a total volume of 10 μl to anneal by the following procedure: denaturation for 5 min at 95°C, cooling to 85°C at -2°C /sec and then cooling from 85°C to 25°C at -0.1°C /sec. The single strand DNA overhangs were filled with T4 DNA polymerase by adding 2.5 μl 10 mM dNTPs mix, 1 μl 10x NEBuffer2, 0.2 μl 100x NEB BSA and 0.5 μl T4 DNA polymerase (NEB) and then incubated at 12°C for 20 min. The resulting double strand DNA was purified using QIAquick PCR purification kit (Qiagen). The gRNAs were transcribed using MEGAshortscript kit (Ambion). gRNAs were treated with DNase, which is included in the kit, and precipitated using lithium-chloride precipitation solution (Ambion). For making Cas9 mRNA to co-inject with gRNAs to zebrafish eggs, we used pCS2-nls-zCas9-nls(Jao et al., 2013) (Addgene) as a template DNA. The template DNA was linearized by NotI (NEB) and purified using QIAquick PCR purification kit. Capped nls-zCas9-nls mRNA was synthesized using mMESSAGE mMACHINE SP6 transcription kit (Ambion) in a volume of 20 μl. The synthesized mRNA was treated with DNase and precipitated with lithium-chloride precipitation solution (Ambion).

To assay and determine the indel mutation by gRNAs, the genome obtained from individual embryos of 1 to 4 dpf or tail clipping for direct sequencing and/or high resolution melt (HRM) analysis was used(Thomas et al., 2014). Embryo or tail fin clips were transferred into 25 to 50 μl of lysis buffer (10 mM Tris-HCl (pH 8.0), 50 mM KCl, 0.3% Tween20, 0.3% NP40, 1 mM EDTA, 0.2 mg/ml Proteinase K (Invitrogen)), and incubated at 55°C for 2 hrs to overnight, followed by the incubation at 98°C for 10 min. For direct sequence, the sequence spanning the gRNA target site was amplified by PCR using 1 μl of the lysed sample as a template, and purified the PCR product. Primers used for the PCR and direct sequencing are listed in Supplementary Table 2. For HRM analysis, we used LightCycler 480 High Resolution Melting Master (Roche) in combination with the LightCycler480 system (Roche). HRM reaction was performed in a volume of 10 μl containing 3.5 mM MgCl_2_ and 0.2 μM each primer by running Gene Scanning 384-II program with the setting of annealing temperature of 60 to 63°C for the amplification and 5 sec hold time and 1°C /s ramp rate at 65°C for high resolution melting. The results were analyzed using the programs of Gene Scanning, Melt Curve Genotyping and Tm calling in LightCycler480 software (Roche). Primers used for HRM analysis are listed in Table S2.

### Rescue experiment

The pTol2(150/250):cmlc2(-210+34):MCS:polyA (pTol2:cmlc2:MCS:pA) plasmid (provided by Dr. Mochizuki) was used to construct pTol2:cmlc2: myl7-P2A-egfp, pTol2:cmlc2: cmlc1-P2A-egfp and pTol2:cmlc2: tnnc1a-P2A-egfp. The myl7, cmlc1 and tnnc1a sequences were amplified by PCR from 4 dpf zebrafish cDNA using following primers; For myl7, forward, 5’-CGC<u>ATCGAT</u>(ClaI)GCCACCATGGCTAGTAAAAAAGCCGCGG-3’, reverse, 5’-CGC<u>GAATTC(EcoRI)</u>AGATTCCTCTTTTTCATCACCATGTGTG-3’, for cmlc1, forward, 5’-CGC<u>ATCGAT</u>(ClaI)GCCACCATGGCACCAAAAAAAGTGGAACC-3’, reverse, 5’-CGC<u>GAATTC(EcoRI)</u>CCCGGAGAGGATGTGCTTGATG-3’, for tnnc1a, forward, 5’-CGC<u>ATCGAT(ClaI)</u>GCCACCATGAACGACATCTACAAAGCAGC-3’, reverse, 5’-CGC<u>AAGCTT(HindIII)</u>TTCCACCCCCTTCATGAACTCC-3’. P2A-egfp fragment amplified from pBluscript:P2A-egfp using primers; forward, 5’-CGC<u>GAATTC(EcoRI)</u>GGAAGCGGAGCTACTAACTTCAGC-3’(for fusion to myl7 and cmlc1) and 5’-CGC<u>AAGCTT(HindIII)</u>GGAAGCGGAGCTACTAACTTCAGC-3’ (for fusion to tnnc1a), reverse, 5’-CGC<u>ACTAGT(SpeI)</u>TTACTTGTACAGCTCGTCCATGCC-3’. ClaI/SpeI-digested pTol2:cmlc2:MCS:pA vector was mixed with one of myl7, cmlc1 and tnnc1a fragments and P2A-egfp fragment to place myl7/cmlc1/tnnc1a-P2A-egfp sequence in the MCS of pTol2:cmlc2:MCS:pA. The pTol2:cmlc2:egfp was constructed by placing egfp sequence in the MCS of pTol2:cmlc2:MCS:pA. 10 ng/μl pTol2:cmlc2(myl7): myl7-P2A-egfp and pTol2:cmlc2:egfp were injected with 25 ng/μl transposase mRNA into one cell stage embryos obtained from myl7^+/+7bp^, cmlc1^+/+29bp^, tnnc1a^+/-5bp^ allele mating.

### Morpholino knockdown

Morpholinos were obtained from Gene Tools, LLC. A translation-blocking morpholino against etv2/etsrp(Pham et al., 2007) (5’-GGTTTTGACAGTGCCTCAGCTCTGC-3’) targeting the -34 to -10 region of the etv2/etsrp sequences was used. The morpholino solution with the concentration of 2 ng/μl was injected to one-cell embryos obtained from *Tg(fli1a:egfp)* and *Tg(gata1:DsRed)* mating. Embryos were harvested on 2 dpf for the analyses.

### DMOG treatment

DMOG (sigma) was diluted in 1x PBS to make 100 mM stock solution. The stock solution was further diluted to prepare the solution of 100 and 125 μM in 1/3 Ringer’s solution containing PTU with 0.1% DMSO. For the control experiment, an equal volume (1 μL) of 1x PBS to the DMOG volume was added to 1/3 Ringer’s solution containing PTU with 0.1% DMSO. About 20 embryos/larvae at appropriate stage were treated in 6 cm petri dish.

### Hypoxia chamber experiment

Hypoxia Incubator Chamber (Billups-Rothenberg) was used to induce body-wide hypoxia. Sibling embryos/larvae in a dish were separated into two groups each in 6 cm dish, containing 10 mL of 1/3 Ringer’s solution containing PTU. One dish was set in hypoxia chamber to flow 5% O_2_/95% N_2_ gas mixture for 5 min with a flow rate of 20 L/min following to the manufacturer’s instruction. Hypoxia chamber was placed in 28.5°C incubator for 24 hrs and then the larvae were harvested.

### Cardiotoxin treatment

Haloperidol (Sigma) was dissolved in DMSO to make the stock solution of 50 mM and 25 mM, respectively. Haloperidol was diluted to 50 μM in 1/3 Ringer’s solution containing PTU. Embryos/larvae were treated with haloperidol for 5 hrs at 28.5°C to 4.5 dpf. Before harvesting, the larvae were assessed under dissection microscope to select those with no heart-beating.

### Atorvastatin treatment

Atorvastatin Calcium Trihydrate (Wako) was dissolved in 100% DMSO at a concentration of 10 mM. Drug was diluted in 0.001% PTU to make the working solution of 2 μM with 0.2% DMSO. Atorvastatin treatment was initiated at 24 hpf and replaced with fresh drug every 24 hrs to 4.5 dpf. To combine atorvastatin treatment and heart ablation, embryos were first treated with atorvastatin to 3 dpf. Then the embryos were soaked in the mixture of 2 μM atorvastatin and 10 mM MTZ for 6 hrs at 3 dpf, followed by the incubation in atorvastatin solution again to 4.5 dpf.

### Data analyses and statistics

For data collection and analysis of qRT-PCR, no statistical methods were used to predetermine sample size. Embryos/larvae subjected to qRT-PCR analysis were blindly collected from the clutch mates. The etv2/etsrp morphants were identified by the reduced expression of *Tg(fli1a:egfp)* reporter and processed for the analyses. For collecting larvae in Figure S4, those expressing GFP widely in heart (more than 70% in heart in appearance under fluorescence stereo microscope) were collected for qRT-PCR analysis. For qRT-PCR data analysis of mutant fish of myl7, cmlc1 and tnnc1a in Fig. 2, values of 2^-dCt^ obtained from two different mutant alleles of each gene were combined to calculate statistics. Because the expression level of lepb gene in wild type embryo/larva was extremely low, a signal of SYBR Green was not detected in our experimental design of qRT-PCR in most of WT. Therefore, in Figs 2 and 6, and Fig. S5, we considered the Ct of lepb as 46, if the signal was not detected. For drug treatment in Fig. S9, embryos/larvae were randomly selected and processed for each treatment group. For the WISH experiments, at least 8 individual fish were processed and examined. Student t-test was performed for statistical analysis. Benjamini-Hochberg procedure was also applied to all multiple sample comparisons (i.e., heatmaps) to correct errors for the multiple tests. A p-value less than 0.05 was considered to be statistically significant (* p<0.05, **p<0.01 and ***p<0.001). The horizontal line represents the mean.

## AUTHOR CONTRIBUTIONS

T.N.S. conceived the idea of the project, designed the overall experiments and *in silico* analyses and supervised the overall research project. N.T. and M.O. contributed to designing the experiments and performed the experiments. S.K. performed the in silico analyses. F.S. and S.E. contributed to the experiments. N.C.C. developed *Tg(cmlc2:mcherry-NTR)* zebrafish line. T.N.S., N.T., M.O., S.K. wrote the manuscript.

### ACKNOWLEDGEMENTS

We thank Y. Oka, T. Ninomiya, T. Kuroda, H. Anabuki for technical assistance, R. Takahashi, E. Kojima for administrative assistance, T. Morie for managing the laboratory’s daily operations. J.H. Neale and S. Kawaoka read the manuscript draft versions and provided critical and constructive comments on the manuscript. We are also grateful to the Sato lab members for advice and discussion throughout the course of this work. M.O. has completed this work as part of her Ph.D. dissertation requirement. M.O. was supported by Grant-in-Aid for JSPS Research Fellow Grant Number 26-10688. This work was supported by JST ERATO Grant Number JPMJER1303 (T.N.S.), JSPS KAKENHI S Grant Number 22229007 (T.N.S.).

## DATA AVAILABILITY

Datasets are available from i-organs.atr.jp.

## SUPPLEMENTARY INFORMATION

Supplementary Information includes 11 figures, 2 tables and 11 movies and can be found with this article online.

## REFERENCES

Aaronson, P.I., Ward, J.P.T., and Connolly, M.J. (2014). The cardiovascular system at a glance, 4th edition edn (United States: Wiley-Blackwell).

Afgan, E., Baker, D., van den Beek, M., Blankenberg, D., Bouvier, D., Cech, M., Chilton, J., Clements, D., Coraor, N., Eberhard, C., et al. (2016). The Galaxy platform for accessible, reproducible and collaborative biomedical analyses: 2016 update. Nucleic acids research 44, W3–W10.

Anderson, J.L., Carten, J.D., and Farber, S.A. (2011). Zebrafish lipid metabolism: from mediating early patterning to the metabolism of dietary fat and cholesterol. Methods in cell biology 101, 111–141.

Butler, J.M., Kobayashi, H., and Rafii, S. (2010a). Instructive role of the vascular niche in promoting tumour growth and tissue repair by angiocrine factors. Nature reviews Cancer 10, 138–146.

Butler, J.M., Nolan, D.J., Vertes, E.L., Varnum-Finney, B., Kobayashi, H., Hooper, A.T., Seandel, M., Shido, K., White, I.A., Kobayashi, M., et al. (2010b). Endothelial cells are essential for the self-renewal and repopulation of Notch-dependent hematopoietic stem cells. Cell stem cell 6, 251–264.

Califano, J.P., and Reinhart-King, C.A. (2010). Exogenous and endogenous force regulation of endothelial cell behavior. J Biomech 43, 79–86.

Camp, J.G., Sekine, K., Gerber, T., Loeffler-Wirth, H., Binder, H., Gac, M., Kanton, S., Kageyama, J.,Damm, G., Seehofer, D., et al. (2017). Multilineage communication regulates human liver bud development from pluripotency. Nature 546, 533–538.

Celik, A., Fuss, S.H., and Korsching, S.I. (2002). Selective targeting of zebrafish olfactory receptor neurons by the endogenous OMP promoter. The European journal of neuroscience 15, 798–806.

Cleaver, O., and Dor, Y. (2012). Vascular instruction of pancreas development. Development 139, 2833–2843.

Craig, M.P., Grajevskaja, V., Liao, H.K., Balciuniene, J., Ekker, S.C., Park, J.S., Essner, J.J., Balciunas, D., and Sumanas, S. (2015). Etv2 and fli1b function together as key regulators of vasculogenesis and angiogenesis. Arteriosclerosis, thrombosis, and vascular biology 35, 865–876.

Crivellato, E., Nico, B., and Ribatti, D. (2007). Contribution of endothelial cells to organogenesis: a modern reappraisal of an old Aristotelian concept. J Anat 211, 415–427.

Curado, S., Anderson, R.M., Jungblut, B., Mumm, J., Schroeter, E., and Stainier, D.Y. (2007).Conditional targeted cell ablation in zebrafish: a new tool for regeneration studies. Developmental dynamics: an official publication of the American Association of Anatomists 236, 1025–1035.

Curado, S., Stainier, D.Y., and Anderson, R.M. (2008). Nitroreductase-mediated cell/tissue ablation in zebrafish: a spatially and temporally controlled ablation method with applications in developmental and regeneration studies. Nature protocols 3, 948–954.

D’Amico, L., Scott, I.C., Jungblut, B., and Stainier, D.Y. (2007). A mutation in zebrafish hmgcr1b reveals a role for isoprenoids in vertebrate heart-tube formation. Current biology: CB 17, 252–259.

Dickover, M.S., Zhang, R., Han, P., and Chi, N.C. (2013). Zebrafish cardiac injury and regeneration models: a noninvasive and invasive in vivo model of cardiac regeneration. Methods in molecular biology 1037, 463–473.

Ding, B.S., Nolan, D.J., Butler, J.M., James, D., Babazadeh, A.O., Rosenwaks, Z., Mittal, V., Kobayashi, H., Shido, K., Lyden, D., et al. (2010). Inductive angiocrine signals from sinusoidal endothelium are required for liver regeneration. Nature 468, 310–315.

Ding, B.S., Nolan, D.J., Guo, P., Babazadeh, A.O., Cao, Z., Rosenwaks, Z., Crystal, R.G., Simons, M., Sato, T.N., Worgall, S., et al. (2011). Endothelial-derived angiocrine signals induce and sustain regenerative lung alveolarization. Cell 147, 539–553.

Fernandes, A.M., Beddows, E., Filippi, A., and Driever, W. (2013). Orthopedia transcription factor otpa and otpb paralogous genes function during dopaminergic and neuroendocrine cell specification in larval zebrafish. PLoS one 8, e75002.

Field, H.A., Dong, P.D., Beis, D., and Stainier, D.Y. (2003a). Formation of the digestive system in zebrafish. II. Pancreas morphogenesis. Developmental biology 261, 197–208.

Field, H.A., Ober, E.A., Roeser, T., and Stainier, D.Y. (2003b). Formation of the digestive system in zebrafish. I. Liver morphogenesis. Developmental biology 253, 279–290.

Gabella, G. (1995). Cardiovascular system. In Gray’s anatomy, L.H. Bannister, M.M. Berry, P. Collins, M. Dyson, J.E. Dussek, and M.W. Ferguson, eds. (London, UK: Churchill Livingstone), pp. 1451–1626.

Gagnon, J.A., Valen, E., Thyme, S.B., Huang, P., Akhmetova, L., Pauli, A., Montague, T.G., Zimmerman, S., Richter, C., and Schier, A.F. (2014). Efficient mutagenesis by Cas9 protein-mediated oligonucleotide insertion and large-scale assessment of single-guide RNAs. PLoS one 9, e98186.

Gerri, C., Marin-Juez, R., Marass, M., Marks, A., Maischein, H.M., and Stainier, D.Y.R. (2017). Hif-1alpha regulates macrophage-endothelial interactions during blood vessel development in zebrafish. Nat Commun 8, 15492.

Gimbrone, M.A., Jr. (1999). Endothelial dysfunction, hemodynamic forces, and atherosclerosis. Thrombosis and haemostasis 82, 722–726.

Ho, S.Y., Thorpe, J.L., Deng, Y., Santana, E., DeRose, R.A., and Farber, S.A. (2004). Lipid metabolism in zebrafish. Methods in cell biology 76, 87–108.

Isogai, S., Lawson, N.D., Torrealday, S., Horiguchi, M., and Weinstein, B.M. (2003). Angiogenic network formation in the developing vertebrate trunk. Development 130, 5281–5290.

Jao, L.E., Wente, S.R., and Chen, W. (2013). Efficient multiplex biallelic zebrafish genome editing using a CRISPR nuclease system. Proceedings of the National Academy of Sciences of the United States of America 110, 13904–13909.

Kajimura, S., Aida, K., and Duan, C. (2005). Insulin-like growth factor-binding protein-1 (IGFBP-1) mediates hypoxia-induced embryonic growth and developmental retardation. Proceedings of the National Academy of Sciences of the United States of America 102, 1240–1245.

Kanai, M., Raz, A., and Goodman, D.S. (1968). Retinol-binding protein: the transport protein for vitamin A in human plasma. The Journal of clinical investigation 47, 2025–2044.

Kimmel, C.B., Ballard, W.W., Kimmel, S.R., Ullmann, B., and Schilling, T.F. (1995). Stages of embryonic development of the zebrafish. Developmental dynamics: an official publication of the American Association of Anatomists 203, 253–310.

Kohn, J.C., Zhou, D.W., Bordeleau, F., Zhou, A.L., Mason, B.N., Mitchell, M.J., King, M.R., and Reinhart-King, C.A. (2015). Cooperative effects of matrix stiffness and fluid shear stress on endothelial cell behavior. Biophysical journal 108, 471–478.

Lammert, E., Cleaver, O., and Melton, D. (2001). Induction of pancreatic differentiation by signals from blood vessels. Science 294, 564–567.

Lammert, E., Cleaver, O., and Melton, D. (2003). Role of endothelial cells in early pancreas and liver development. Mech Dev 120, 59–64.

Langmead, B., and Salzberg, S.L. (2012). Fast gapped-read alignment with Bowtie 2. Nature methods 9, 357–359.

Lawson, N.D., Vogel, A.M., and Weinstein, B.M. (2002). sonic hedgehog and vascular endothelial growth factor act upstream of the Notch pathway during arterial endothelial differentiation. Developmental cell 3, 127–136.

Lawson, N.D., and Weinstein, B.M. (2002). In vivo imaging of embryonic vascular development using transgenic zebrafish. Developmental biology 248, 307–318.

Li, E., and Norris, A.W. (1996). Structure/function of cytoplasmic vitamin A-binding proteins. Annu Rev Nutr 16, 205–234.

Liao, W., Bisgrove, B.W., Sawyer, H., Hug, B., Bell, B., Peters, K., Grunwald, D.J., and Stainier, D.Y. (1997). >The zebrafish gene cloche acts upstream of a flk-1 homologue to regulate endothelial cell differentiation. Development 124, 381–389.

Love, M.I., Huber, W., and Anders, S. (2014). Moderated estimation of fold change and dispersion for RNA-seq data with DESeq2. Genome biology 15, 550.

Lu, L., Chen, Y., Wang, Z., Li, X., Chen, W., Tao, Z., Shen, J., Tian, Y., Wang, D., Li, G., et al. (2015).The goose genome sequence leads to insights into the evolution of waterfowl and susceptibility to fatty liver. Genome biology 16, 89.

Manchenkov, T., Pasillas, M.P., Haddad, G.G., and Imam, F.B. (2015). Novel Genes Critical for Hypoxic Preconditioning in Zebrafish Are Regulators of Insulin and Glucose Metabolism. G3(Bethesda) 5, 1107–1116.

Mazein, A., Watterson, S., Hsieh, W.Y., Griffiths, W.J., and Ghazal, P. (2013). A comprehensive machine-readable view of the mammalian cholesterol biosynthesis pathway. Biochemical pharmacology 86, 56–66.

Milan, D.J., Jones, I.L., Ellinor, P.T., and MacRae, C.A. (2006). In vivo recording of adult zebrafish electrocardiogram and assessment of drug-induced QT prolongation. American journal of physiology Heart and circulatory physiology 291, H269–273.

Mukouyama, Y.S., Shin, D., Britsch, S., Taniguchi, M., and Anderson, D.J. (2002). Sensory nerves determine the pattern of arterial differentiation and blood vessel branching in the skin. Cell 109, 693–705.

Myers, M.G., Jr., and Olson, D.P. (2012). Central nervous system control of metabolism. Nature 491,357–363.

Nellore, A., Jaffe, A.E., Fortin, J.P., Alquicira-Hernandez, J., Collado-Torres, L., Wang, S., Phillips Iii, R.A., Karbhari, N., Hansen, K.D., Langmead, B., et al. (2016). Human splicing diversity and the extent of unannotated splice junctions across human RNA-seq samples on the Sequence Read Archive.Genome biology 17, 266.

Noble, M. (2002). The cardiovascular system in health and disease (London, UK: Imperial College Press).

Okabe, K., Kobayashi, S., Yamada, T., Kurihara, T., Tai-Nagara, I., Miyamoto, T., Mukouyama, Y.S., Sato, T.N., Suda, T., Ema, M., et al. (2014). Neurons limit angiogenesis by titrating VEGF in retina.Cell 159, 584–596.

Paton, C.M., and Ntambi, J.M. (2009). Biochemical and physiological function of stearoyl-CoA desaturase. American journal of physiology Endocrinology and metabolism 297, E28–37.

Pham, V.N., Lawson, N.D., Mugford, J.W., Dye, L., Castranova, D., Lo, B., and Weinstein, B.M. (2007). Combinatorial function of ETS transcription factors in the developing vasculature. Developmental biology 303, 772–783.

Rabbany, S.Y., Ding, B.S., Larroche, C., and Rafii, S. (2013). Mechanosensory pathways in angiocrine mediated tissue regeneration (Berlin: Springer).

Rafii, S., Butler, J.M., and Ding, B.S. (2016). Angiocrine functions of organ-specific endothelial cells.Nature 529, 316–325.

Reinhart-King, C.A., Fujiwara, K., and Berk, B.C. (2008). Physiologic stress-mediated signaling in the endothelium. Methods in enzymology 443, 25–44.

Reischauer, S., Stone, O.A., Villasenor, A., Chi, N., Jin, S.W., Martin, M., Lee, M.T., Fukuda, N., Marass, M., Witty, A., et al. (2016). Cloche is a bHLH-PAS transcription factor that drives haemato-vascular specification. Nature 535, 294–298.

Santhakumar, K., Judson, E.C., Elks, P.M., McKee, S., Elworthy, S., van Rooijen, E., Walmsley, S.S.,Renshaw, S.A., Cross, S.S., and van Eeden, F.J. (2012). A zebrafish model to study and therapeutically manipulate hypoxia signaling in tumorigenesis. Cancer research 72, 4017–4027.

Sato, T.N. (2013). Mechanical and chemical regulation of arterial and veous specification (Berlin:Springer).

Semenza, G.L. (2011). Oxygen sensing, homeostasis, and disease. The New England journal of medicine 365, 537–547.

Semenza, G.L. (2014). Oxygen sensing, hypoxia-inducible factors, and disease pathophysiology. Annual review of pathology 9, 47–71.

Serluca, F.C., Drummond, I.A., and Fishman, M.C. (2002). Endothelial signaling in kidney morphogenesis: a role for hemodynamic forces. Current biology: CB 12, 492–497.

Simon, M.C., and Keith, B. (2008). The role of oxygen availability in embryonic development and stem cell function. Nature reviews Molecular cell biology 9, 285–296.

Sumanas, S., and Lin, S. (2006). Ets1-related protein is a key regulator of vasculogenesis in zebrafish.PLoS Biol 4, e10.

Talavera-Adame, D., and Dafoe, D.C. (2015). Endothelium-derived essential signals involved in pancreas organogenesis. World J Exp Med 5, 40–49.

Trapnell, C., Williams, B.A., Pertea, G., Mortazavi, A., Kwan, G., van Baren, M.J., Salzberg, S.L.,Wold, B.J., and Pachter, L. (2010). Transcript assembly and quantification by RNA-Seq reveals unannotated transcripts and isoform switching during cell differentiation. Nature biotechnology 28, 511–515.

Traver, D., Paw, B.H., Poss, K.D., Penberthy, W.T., Lin, S., and Zon, L.I. (2003). Transplantation and in vivo imaging of multilineage engraftment in zebrafish bloodless mutants. Nature immunology 4, 1238–1246.

Veldman, M.B., and Lin, S. (2012). Etsrp/Etv2 is directly regulated by Foxc1a/b in the zebrafish angioblast. Circulation research 110, 220–229.

Visconti, R.P., Richardson, C.D., and Sato, T.N. (2002). Orchestration of angiogenesis and arteriovenous contribution by angiopoietins and vascular endothelial growth factor) VEGF). Proceedings of the National Academy of Sciences of the United States of America 99, 8219–8224.

Wang, H.U., Chen, Z.F., and Anderson, D.J. (1998). Molecular distinction and angiogenic interaction between embryonic arteries and veins revealed by ephrin-B2 and its receptor Eph-B4. Cell 93, 741–753.

Weinstein, B.M. (2005). Vessels and nerves: marching to the same tune. Cell 120, 299–302.

Wu, Y.C., Chang, C.Y., Kao, A., Hsi, B., Lee, S.H., Chen, Y.H., and Wang, I.J. (2015). Hypoxia-induced retinal neovascularization in zebrafish embryos: a potential model of retinopathy of prematurity. PloS one 10, e0126750.

Yoshida, T., Ito, A., Matsuda, N., and Mishina, M. (2002). Regulation by protein kinase A switching of axonal pathfinding of zebrafish olfactory sensory neurons through the olfactory placode-olfactory bulb boundary. The Journal of neuroscience: the official journal of the Society for Neuroscience 22,4964–4972.

Yoshida, T., and Mishina, M. (2003). Neuron-specific gene manipulations to transparent zebrafish embryos. Methods in cell science: an official journal of the Society for In Vitro Biology 25, 15–23.

Zhu, D., Fang, Y., Gao, K., Shen, J., Zhong, T.P., and Li, F. (2017). Vegfa Impacts Early Myocardium Development in Zebrafish. Int J Mol Sci 18.

